# Phosphoproteomics identifies PI3K inhibitor-selective adaptive responses in pancreatic cancer cell therapy and resistance

**DOI:** 10.1101/2020.10.08.313833

**Authors:** C Cintas, T Douche, Z Dantes, E Mouton-Barbosa, MP Bousquet, C Cayron, N Therville, F Pont, F Ramos-Delgado, C Guyon, B Garmy-Susini, P Cappello, O Burlet-Schiltz, E Hirsch, A Gomez-Brouchet, B Thibault, M Reichert, J Guillermet-Guibert

## Abstract

The PI3K pathway is highly active in human cancers. The four class I isoforms of PI3K are activated by distinct mechanisms leading to a common downstream signaling. Their downstream redundancy is thought to be responsible for treatment failures of PI3K inhibitors. We challenged this concept, by mapping the differential phosphoproteome evolution in response to PI3K inhibitors with different isoform selectivity patterns in pancreatic cancer, a disease currently without effective therapy. In this cancer, the PI3K signal was shown to control cell proliferation. We compared the effects of LY294002 that inhibit with equal potency all class I isoenzymes and downstream mTOR with the action of inhibitors with higher isoform-selectivity towards PI3Kα, PI3Kβ or PI3Kγ (namely A66, TGX-221 and AS-252424). A bioinformatics global pathway analysis of phosphoproteomics data allowed us to identify common and specific signals activated by PI3K inhibitors supported by the biological data. AS-252424 was the most effective treatment and induced apoptotic pathway activation as well as the highest changes in global phosphorylation-regulated cell signal. However, AS-252424 treatment induced re-activation of Akt, therefore decreasing the treatment outcome on cell survival. Reversely, AS-252424 and A66 combination treatment prevented p-Akt reactivation and led to synergistic action in cell lines and patient organoids. The combination of clinically approved α-selective BYL-719 with γ-selective IPI-549 was more efficient than single molecule treatment on xenograft growth. Mapping unique adaptive signaling responses to isoform-selective PI3K inhibition will help to design better combinative treatments that prevent the induction of selective compensatory signals.

**Graphical abstract:** 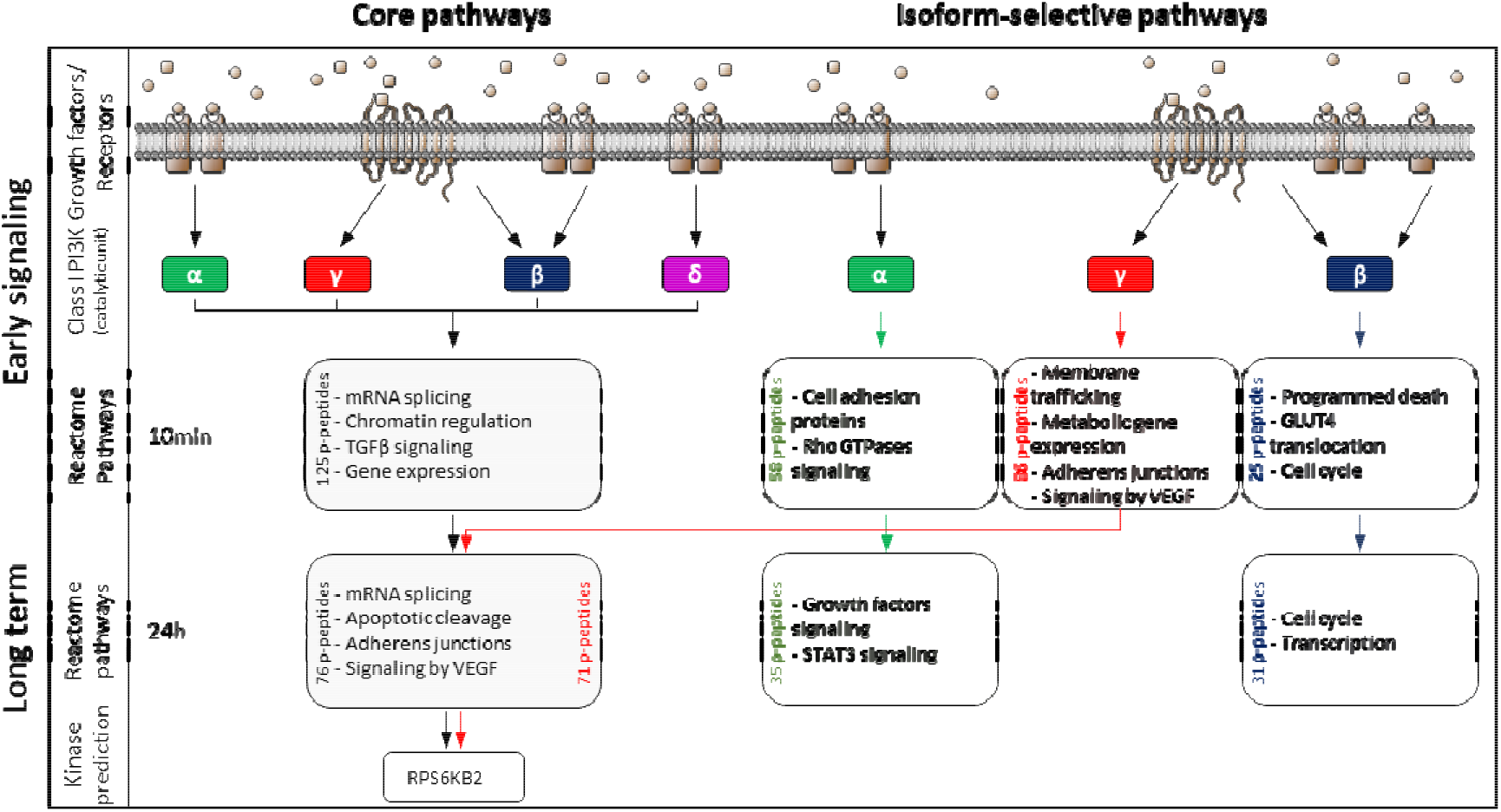

**Significance:** Half of all human cancers show increased PI3K activity but the mutational pattern of the PI3K pathway is not sufficient to predict sensitivity to PI3K inhibitors. By identifying for the first time specific signaling induced by PI3K inhibitors with different isoform-selectivity patterns, we provide insight in how to handle heterogeneity of PI3K expression in tumor samples for the choice of available PI3K-targetting drugs. Our work provides a roadmap to target the PI3K pathway, balancing the use of isoform-selective PI3K inhibitors with personalized information extending beyond the specific mutational status.

## Introduction

In pathophysiological signaling, biochemical and biomechanical cues are integrated and regulated at long-term. Similarly, it is expected that inhibition of signaling pathways by targeted therapies towards one signal transduction enzyme also induces an adaptation of the entire signaling network.

Class I PI3Ks are crucial signal transduction enzymes. After acute stimulation, PI3K phosphorylates the lipid second messenger phosphatidylinositol 4,5-biphosphate into PI-3,4,5-triphosphate (PIP_3_) at the plasma membrane, further activating the protein kinases Akt and mTOR, and regulating major cell biology events such as cell proliferation, cell survival and protein synthesis. PI3K is one of the most altered pathways in cancers, and presents 4 different isoforms encoded by 4 different genes (*1*, *2*). Due to the regulation of fundamental cellular processes by PI3K/Akt/mTOR, this signaling axis is an excellent therapeutic target in cancer which is underscored by the number of molecules tested currently in clinical trials (*3*). While class I PI3K isoform specificity is well described and accepted in physiology (for review: (*1*), examples: (*4*–*7*)), potential benefits of isoform-selective targeting in solid cancer were recently shown in breast cancers driven by oncogenic PI3Kα (*8*) but are still not yet approved in other genetic context promoting PI3K signaling (*3*). Besides, there are other effectors downstream PI3Ks than Akt/mTOR (*9*, *10*). Such other downstream signaling routes are possibly contributing to the isoform-specific *in vivo* role of mammalian PI3Ks. Although acquired cross-activation mechanisms between PI3K isoforms upon their unique and selective pharmacological or genetic inhibition in the context of specific mutational landscapes (e.g. oncogenic *PIK3CA*, mutant *PTEN*) has been described (*11*–*13*), a specific large scale cell signal adaptation to such treatment is unknown, in particular, in the context of non-mutated PI3Ks and/or Kras mutation. This genetic context is found in pancreatic ductal adenocarcinoma (PDAC). It is also unclear whether differing inhibition of each PI3K due to distinct PI3K inhibitor selectivity or due to different isoform expression could favor selective feedback mechanisms; the latter could be an unprecedented explanation of the therapeutic failure of PI3K inhibitors in unselected solid tumours.

New strategies are needed for the cure of pancreatic cancer (PDAC) patients, due to dramatic lethality rate of this disease. PI3K signaling as assessed by Akt phosphorylation or by PI3K/Akt/mTOR gene signature is increased and associated with poor prognosis (*14*, *15*). This increase in PI3K signaling is mostly due to Kras mutation engaging PI3Kα and possibly PI3Kγ (*16*–*19*). PI3K signaling downstream Kras is amplified by other signaling cues (*20*) and such amplification of Kras-PI3K coupling is critically involved in PDAC poor prognosis (*21*). Targeting PI3K is expected to have a clinical action in these patients (clinical trials pending as reviewed in (*22*) and (*23*)). However, prior knowledge of cancer cell adaptation to the inhibition of upstream class I PI3K signaling would be necessary to develop efficient anti-PI3K therapeutic strategy in pancreatic disease (*22*). The purpose of the study is to compare the pancreatic cancer cell phosphoproteome in response to inhibitors with different selectivity and off-targets profiles, as an exploratory experiment to find first evidence of isoform-selective pathways and adaptive responses.

Therefore, we analyzed adaptive signals involved in response to inhibitors with distinct PI3K-isoform selectivity in a human pancreatic cell line using a stable isotope labeling with amino acids in cell culture (SILAC)-based quantitative phosphoproteomics approach in a comprehensive way. By enrichment of the phosphoproteome followed by data mining, we demonstrate that, despite a common core pathway regulated by all inhibitors, the different PI3K inhibitors instruct specific, non-redundant signaling pathways linked to PI3K signaling. These data might help to better design therapeutic strategies in this dismal disease, including combination therapies against multiple isoforms of PI3K.

## Results

### PI3K inhibitors induce different adaptive phospho-proteome responses

In pancreatic cancer, PI3K signaling is associated with a poor prognosis (*14*, *15*). The analysis of 11 pancreatic cancer samples compared to normal adjacent tissue showed increase Akt expression and a significant increase of S473 and T308 Akt phosphorylation by western blot (WB) (9 out of 11 patients) (**Figure 1A**, **Supplementary Figure 1A**). However, the observed increase of Akt phosphorylation was not always associated with a significant increase in the phosphorylation levels of canonical downstream targets, such as pPRAS40 or pS6K (**Figure 1A**, **right**), emphasizing the heterogeneity of signaling targets downstream PI3Ks in a clinical setting.

**Figure 1:**
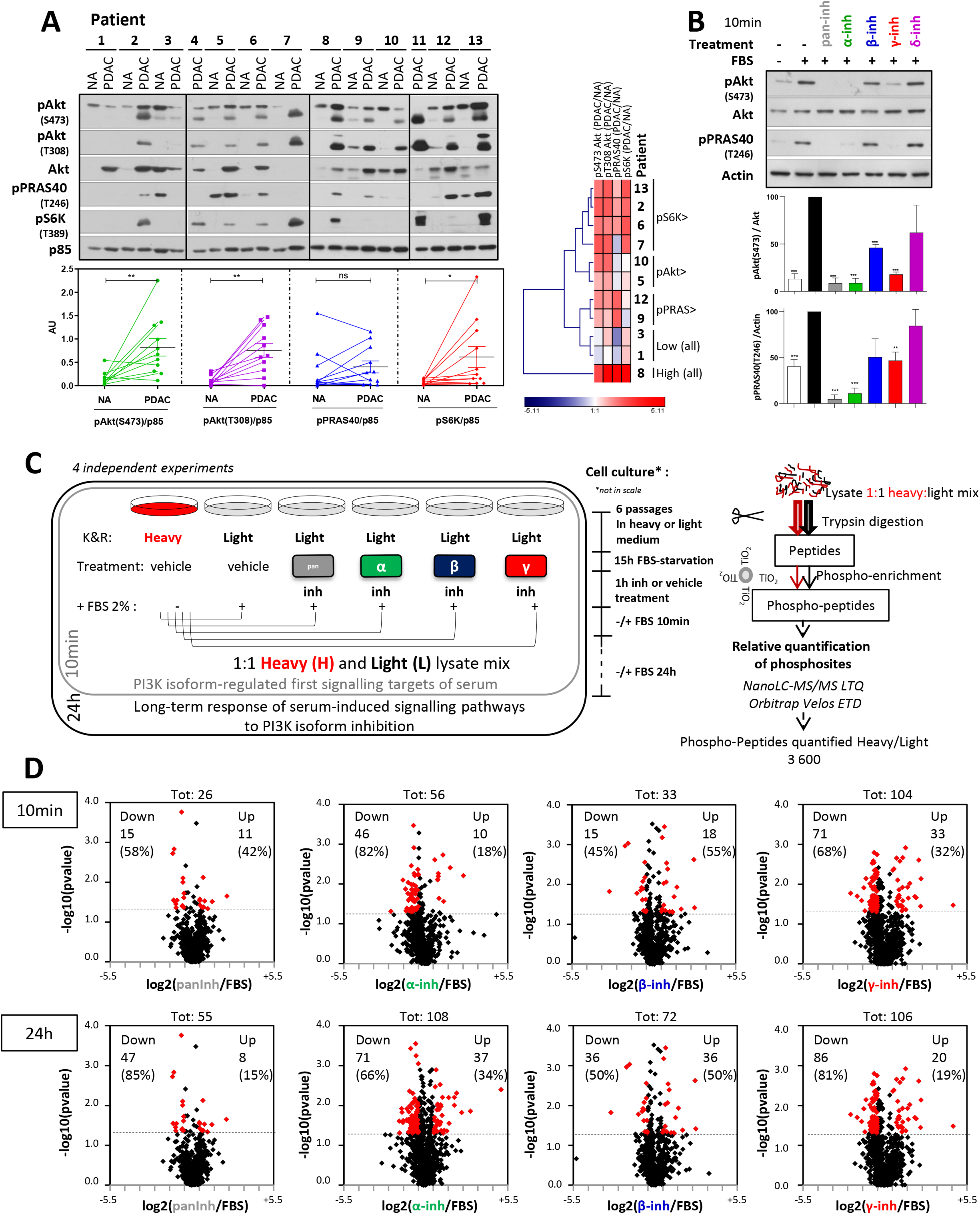
Level of PI3K/Akt pathway activation in pancreatic cancer and strategy for the identification of core redundant PI3K-regulated signaling networks versus isoform-selective signaling networks. (A) Frozen samples of normal adjacent pancreas (NA, n=11 patients) and pancreatic cancer (PDAC, n=13 patients) samples were lysed. Phospho-effector abundance of PI3K/Akt/mTOR pathway was assessed by western blot using indicated antibodies, in samples comprising >30% of epithelial cells; quantification was performed only when the normal adjacent tissue was available (n=11 patients). Hierarchical clustering is shown using log (fold-change PDAC vs. normal adjacent). (B) Capan-1 pancreatic cancer cells were pre-treated with pan-PI3K, isoform-selective inhibitors (same conditions) or their accordingly diluted vehicle (DMSO) during 1h, then stimulated or not for 10min with 2% FBS. After 10min cells were lysed and phospho-effector abundance of PI3K/Akt/mTOR pathway was assessed by western blot using indicated antibodies and quantified (n≥4 independent biological replicates). Total Akt and Actin is used as a loading control. (C, D) Scheme of SILAC strategy to identify common or pan-PI3K and isoform-selective phosphorylation-regulated signaling pathways. Light amino acid-labelled and heavy amino acid-labelled Capan-1 cells were pre-treated respectively with 10μM LY-294002 (pan-inh), 5μM A66 (PI3Kα inhibitor named α-inh), 500nM TGX-221 (PI3Kβ inhibitor named β-inh), 5μM AS-252424 (PI3Kγ inhibitor named γ-inh) or their accordingly diluted vehicle (DMSO) during 1h, then stimulated or not for 10min or 24h with 2% dialysed FBS. After 10min and 24h of FBS stimulation, heavy cells and light cells were lysed. For each biological replicate, light and heavy lysates were mixed at a 1:1 ratio, subjected to tryptic digestion and enrichment or not in phosphopeptides with TiO2 beads. Peptides were processed by mass spectrometry (Velos LTQ Orbitrap). Four biological replicates were sampled for each time point; light condition varies in the biological replicate while heavy vehicle condition is a technical replicate that is added in each run. Significantly increased/decreased phospho-peptides ratios (fold change = 2) for each condition are represented in D.

The human pancreatic cancer cell line Capan-1 represents common genetic alterations found in PDAC including *KRAS, TP53 and SMAD4* mutations (*24*), expresses all four PI3K isoforms (*16*, *25*) and displays a proliferation rate similar to other human pancreatic cell lines (as verified in **Supplementary Figure 1B**). Amongst the four PI3K isoforms responsible for the production of PIP_3_ and Akt activation, we and others have identified PI3Kα and PI3Kγ to be involved in pancreatic carcinogenesis (*16*, *17*, *26*–*28*). Besides oncogenic Kras-driven activation of PI3K (*20*), stimulation by fetal bovine serum (FBS) induces the activation of receptor tyrosine kinases (RTKs) and G protein-coupled receptors (GPCRs) that increases pAkt level (*7*). Short-term (10min) FBS stimulation induced a significant activation of class I PI3Ks as assessed by the phosphorylation of Akt and a known downstream effector, PRAS40 (**Figure 1B**). We used a pan-PI3K/mTOR-targeting inhibitor that inhibits all PI3K isoforms at equal potency [LY-294002 from here on named pan-inh]. Pan-inh completely abolished pAkt and pPRAS40 (**Figure 1B**). Isoform-selective drugs targeting either PI3Kα (A66 [5μM], α-inh (*29*)), PI3Kβ (TGX-221 [0.5μM], β-inh (*7*, *30*–*32*)) and PI3Kγ (AS-252424 [5μM], γ-inh (*33*)), but not of PI3Kδ inhibitor (IC-87114 [5μM], δ-inh), inhibited pS473Akt as well as pPRAS40 levels significantly after 10min of FBS-stimulation (**Figure 1B**). PI3K inhibitors are still effective to inhibit pAkt when diluted in cell medium 24h prior to treatment *in vitro* demonstrating the stability of the compounds at long-term (**Supplementary Figure 1C**).

To explore the possibility of isoform-selective downstream pathways, we devised an exploratory strategy to identify adaptive response to PI3K inhibitors with varying selectivity and off-targets effects in a comprehensive fashion by defining phospho-site regulated signaling pathways in Capan-1 cell line (**Figure 1C**). To quantify subtle differences between these conditions targeting the same enzymatic activity, we chose a SILAC-based quantitative approach combined to a phospho-peptide enrichment by TiO2 allowing a robust S/T/Y phosphorylation quantification of thousands of proteins. We devised a super-SILAC approach (*34*, *35*), in which we compared all the unlabeled treatment conditions to vehicle control, with the use of SILAC-labeled cells as a spike-in standard for accurate quantification of unlabeled samples (**Figure 1C**, **Supplementary Table 1**). Incorporation of heavy isotopes was verified by mass spectrometry after 6 passages (**Supplementary Figures 2A, 2B**); and the metabolic labelling did not change the proliferation and morphological properties of Capan-1 cell line (**Supplementary Figures 2C, 2D**). All validating steps are detailed in **Supplemental Material & Methods** and **Supplementary Figures 2**. In all conditions combined, 3600 heavy/light phospho-peptides were detected and quantified (**Figure 1C**). Amongst these, 83% serine-sites (S), 16% threonine-sites (T), 1% tyrosine-sites (Y) were phosphorylated (**Supplementary Figure 2F**). These percentages were unchanged upon PI3K inhibition (**Supplementary Figure 2G**). Short-term (10min)- and long-term (24h)-serum stimulation induced modifications (increased and decreased) of 557 phospho-peptides (28%) and 619 phospho-peptides (32%), respectively (**Supplementary Figures 2H, 2I**).

Overall, levels of a known PI3K target PRAS40 were changed in similar manner albeit with a slightly lower dynamic range when quantified by MS-based proteomics analysis compared to western blot results (**Supplementary Figures 2J** vs. **Figure 1B**).

We next identified phospho-peptides with significant altered levels in each condition in an unbiased fashion. PI3K inhibitors induced more phosphorylation level changes after 24h of treatment than after 10min. Interestingly, γ-inh led to the most significant changes in numbers of significantly modified phosphopeptides compared to FBS as soon as 10min of treatment, and global equivalent phosphoprotein level changes at 24h compared to α-inh (**Figure 1D**). These strong phosphoprotein modulations by γ-inh and α-inh after 24h of treatment suggest signaling engagement of these two PI3K isoforms in PDAC cells, in addition to their involvement in pancreatic carcinogenesis (*16*, *17*, *26*–*28*).

The number of significantly increased p-peptides in γ-inh condition (and to slighter extent of pan-inh) was higher at short time compared to α-inh but decreased at 24h, possibly suggesting an early induction of feedback control with these two PI3K inhibitors. Reversely, α-inh treatment led to increased p-peptides at 24h. β-inh treatment led to a balanced increase/decrease of p-peptides at both times (**Figure 1D**). Early upregulated signaling is expected to reduce the efficiency of PI3Kγ inhibitors.

### Global pathway analysis of phosphoproteome upon PI3K inhibition shows selective changes associated with different isoform-specificity

We next analyzed dynamic changes in the phosphoproteome upon PI3K isoform-specific inhibition over time (10min vs. 24h treatment) using global pathway analysis 1- to identify selectivity in regulating biological functions by each inhibitor, and 2- visualize entire signal network rewiring upon PI3K inhibition pressure.

A principal component analysis (PCA) of the phosphoproteome at 10min demonstrated that serum-stimulation alone, α-inh and β-inh conditions clustered together while pan-inh and γ-inh conditions separated from the cluster (**Figure 2A**, **left**). After 24h of treatment, however, all inhibitor treatments separated from the FBS condition, highlighting the time necessary to induce significant PI3K inhibitor-selective changes in signaling networks upon PI3K inhibition, particularly for PI3K isoform-specific inhibition (**Figure 2A**, **right**). Surprisingly, inhibition with LY (pan-inh) clustered closely together with the β-inh, while γ- and α-inh conditions led to distinct phosphopeptide modifications, in line with the known selective roles of these isoforms in PDAC (*17*, *25*, *26*).

**Figure 2:**
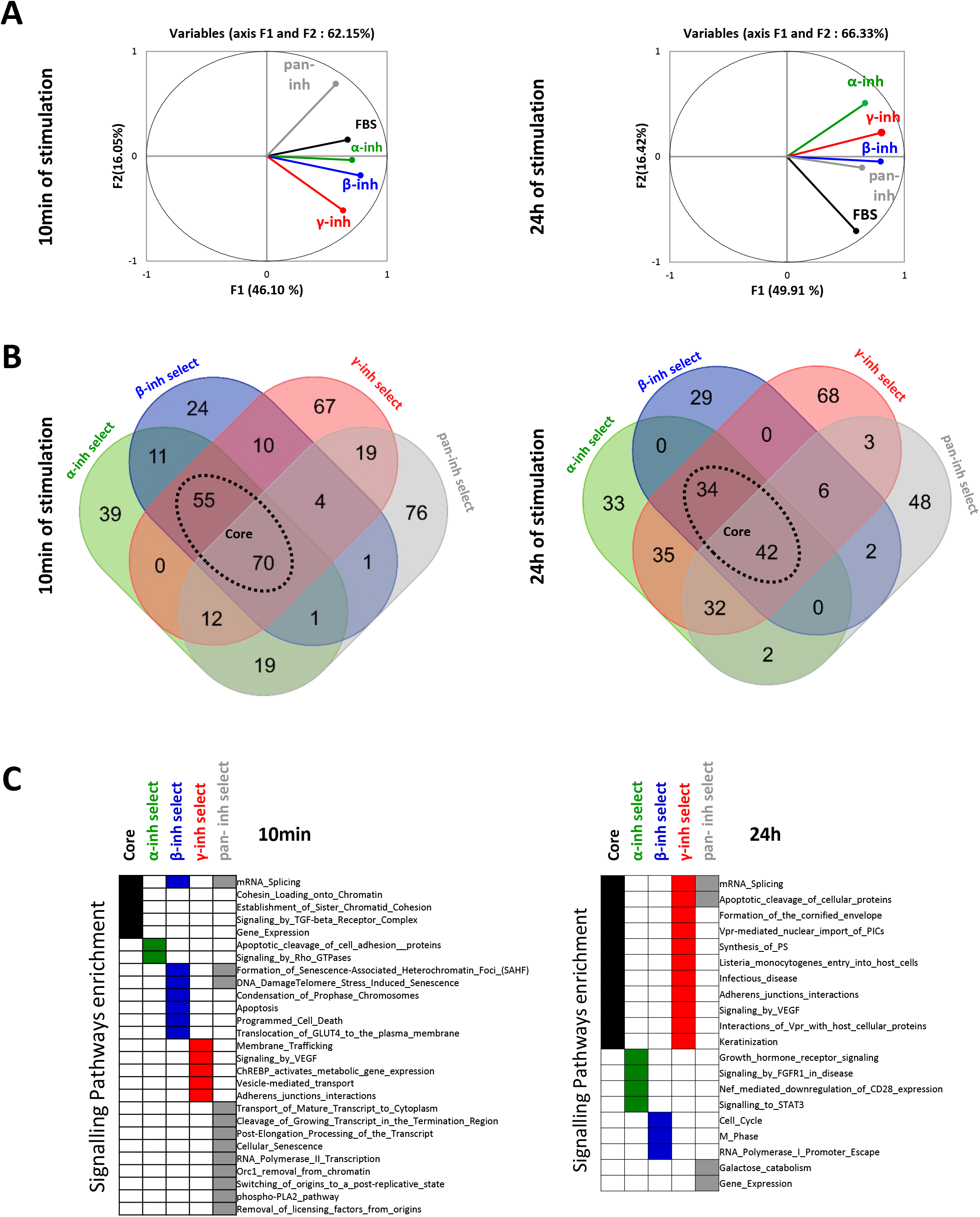
Class I PI3K isoforms regulate distinct signaling pathways in pancreatic cancer cells. (A) Global variation of phosphoproteome under FBS stimulation with or without PI3K inhibitor treatment in Capan-1 at 10min and 24h. Principal component analysis (PCA) analysis is shown. n=4 independent biological replicates. (B) Representation of targets specifically modified by pan-PI3K, PI3Kα, PI3Kβ and PI3Kγ inhibition. Dashed lines explain how core phosphopeptides are defined. A Venn diagram of modified phosphopeptides in at least one condition at each time point is shown. n=4 independent biological replicates. (C) Enrichments of biological pathways found in the list of core, pan-, isoform-selective phosphoproteins at 10min and 24h were performed with AutoCompare software (pvalue ≤ 0.01). n=4 independent biological replicates.

Next, we identified inhibitor-selectivity in regulated phosphoproteomes at both time points using hierarchical clustering (**Supplemental Material and Methods, Supplementary Table 1, 2, 3**). By overlapping phosphopeptides with altered levels in each condition, we were able to identify shared (core) phosphopeptides, inhibitor-selective phosphopeptides indicated as either A66-selective (α-inh -select), TGX-221-selective (β-inh -select) or AS252424 -selective (γ-inh select), as well as pan/mTOR-selective (pan-inh-select) phosphopeptides (**Figures 2B, 2C**, **Supplementary Figure 3**). The finding that there is a common “core” of phosphopeptides regulated by all PI3K inhibitors was underscored by STRING analyses of inhibitor-selective and common phosphopeptides **(Supplementary Figures 3A, 3B, Supplementary Table 1, 2, 3**). This core network presented a strong connectivity at 10min of treatment.

Our data also demonstrate that each inhibitor induced distinct changes of the phosphorylation-regulated proteome already at 10min of treatment (**Figure 2A, 2B)**, despite having similar effects on the phosphorylation of Akt and PRAS40 (**Figure 1B**). Inhibitor-selective as well as core phosphopeptides showed distinct pathway enrichments (Reactome), cellular components and molecular functions (Gene Ontology) at the 10min and 24h time points (**Figure 2C**, **Supplementary Figure 3**). At 10min, common core downstream signaling includes 115 phosphoproteins implicated in mRNA splicing, chromatin regulation, TGFβ signaling as well as gene transcription and transcript processing pathways (ZE Pvalue ≤0.05), such as EIF3G, EIF2S2, EIF4G1 (**Figures 2C**, **Supplementary Table 2**). Interestingly, at 24h, γ-inh-selective phosphopeptides were enriched in similar functions and cellular components as those changed by all PI3Ks/mTOR (core and pan-inh select), with targets presenting a strong protein-protein interactivity; those included CTNND1, JUP, SRRM2 proteins (**Figure 2C**, **Supplementary Figures 3F-J, Supplementary Table 2**). Apart from the core pathways, α-inh-selective and β-inh-selective phosphopeptides were enriched in different pathways at both time points. Specifically, α-inh modulated phosphopeptides regulating Rho GTPases signaling, RTK and cytokine signaling (including RACGAP1, IRS2, STAT3), whereas β-inh regulated phosphopeptides directing mitotic control including targets such as RB1 (**Figure 2C**, **Supplementary 3D, E,G-J, Supplementary Table 2**).

To estimate on-target and off-target effects of inhibitors, we used STRING connectivity analysis and showed that inhibitor selective targets were associated and connected with PI3K isoforms, except for TGX-221 treatment at 24h (**Supplementary Figures 3E, Supplementary Table 1, 2, 3**). The proteins involved were either off-targets of TGX-221 (β-inh) or were unknown targets of PI3Kβ. These data show that PI3K inhibition with AS252424 (γ-inh) regulates similar processes to pan-inh and, at the same time, controls additional core networks exceeding the effect of pan-inh on signal network as shown by the number of non-selective and selective phosphopeptides modified by γ-inh treatment (**Figure 2B**).

These findings suggest that inhibiting strongly PI3Kγ might be the most effective therapeutic strategy in this cell line to inhibit the core PI3K signaling in order to target essential growth and survival pathways in pancreatic cancer.

### Pancreatic cancer cells present increased levels of p110γ expression in patients

We next analyzed the expression levels of PI3K catalytic subunits that constitute the four class I isoforms (α, β, γ and δ) in tumor cell-enriched human pancreatic cancer samples compared to normal adjacent pancreas by Western Blot. We observed that p110β levels were increased only in 5 patients, while p110α, p110γ and p110δ protein levels were significantly increased (in all but one patient) (**Figure 3A**). To further verify these findings in a pure population of cancer cells, we analyzed PI3K isoforms protein and mRNA levels in human (n=4) and murine (n=7) cell lines (derived from genetically modified mouse models of PDAC) (**Figure 3B-E**). The myeloid cell line, MOLM-14, which is known to express p110γ in high abundancy served as positive control.

**Figure 3:**
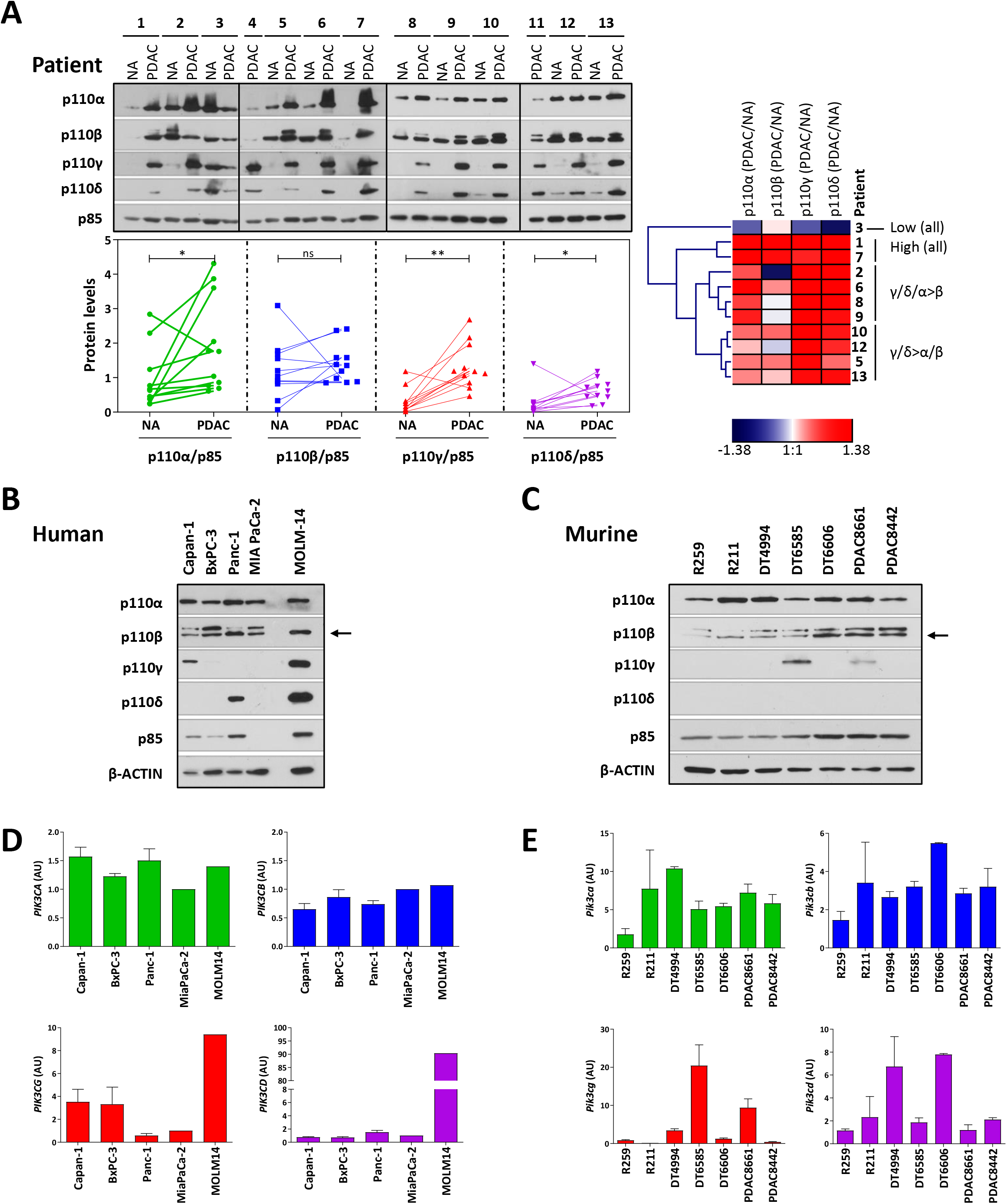
PI3Kγ is overexpressed albeit at low levels in pancreatic cancer tumor cells. (A) Frozen samples of normal adjacent pancreas (NA, n=11 patients) and pancreatic cancer (PDAC, n=13 patients) samples were lysed. PI3K isoforms proteins pattern of expression was assessed by western blot using indicated antibodies, in samples comprising >30% of epithelial cells, and quantified when the adjacent tissue was available. Hierarchical clustering is shown using log (fold-change PDAC vs. normal adjacent). (B-C) Human and murine pancreatic cells were lysed. PI3K isoform mRNA and protein pattern of expression was evaluated by RT-qPCR and WB. Mean ± SEM is shown, WB: n=3 human n=1 murine, qPCR: n=3 human, n=2 murine, independent biological replicates.

In contrast to tumor cell-enriched human pancreatic cancer samples, p110γ was only found detected in one out of four human pancreatic cancer cell lines and in two out of seven murine pancreatic cancer cell lines by Western Blot (**Figure 3B, 3C**). Similarly, on a transcript level, two out of four human PDAC cell lines and three out of seven murine PDAC cell lines showed increase in p110γ mRNA expression by RT-qPCR (**Figure 3D, 3E**). This data indicates that p110γ is overexpressed in patient-derived tumor cell-enriched PDAC samples and to a lesser extent in PDAC cells *in vitro,* suggesting a stronger role for p110γ *in vivo* and in patients, compared to cell culture conditions.

### Pancreatic cancer cells are sensitive to PI3Kγ inhibition *in vitro* and *in vivo*

We next confirmed that genetic inactivation of PI3Kγ using a full knock-out approach almost completely abolished cancer formation from precancer lesions (PanIN) in a mutated KRAS and p53 background (**Figure 4A**). Similarly, despite the low expression levels of p110γ *in vitro*, decreased expression of p110γ using a shRNA specific for p110γ reduced the clonogenicity of Capan-1 cell line (**Figure 4B**). Of note, we did observe decreased PI3Kα expression upon p110γ knockdown by shRNA in one pool of cells, but that led to similar effects that the other shp110γ pool of cells.

**Figure 4:**
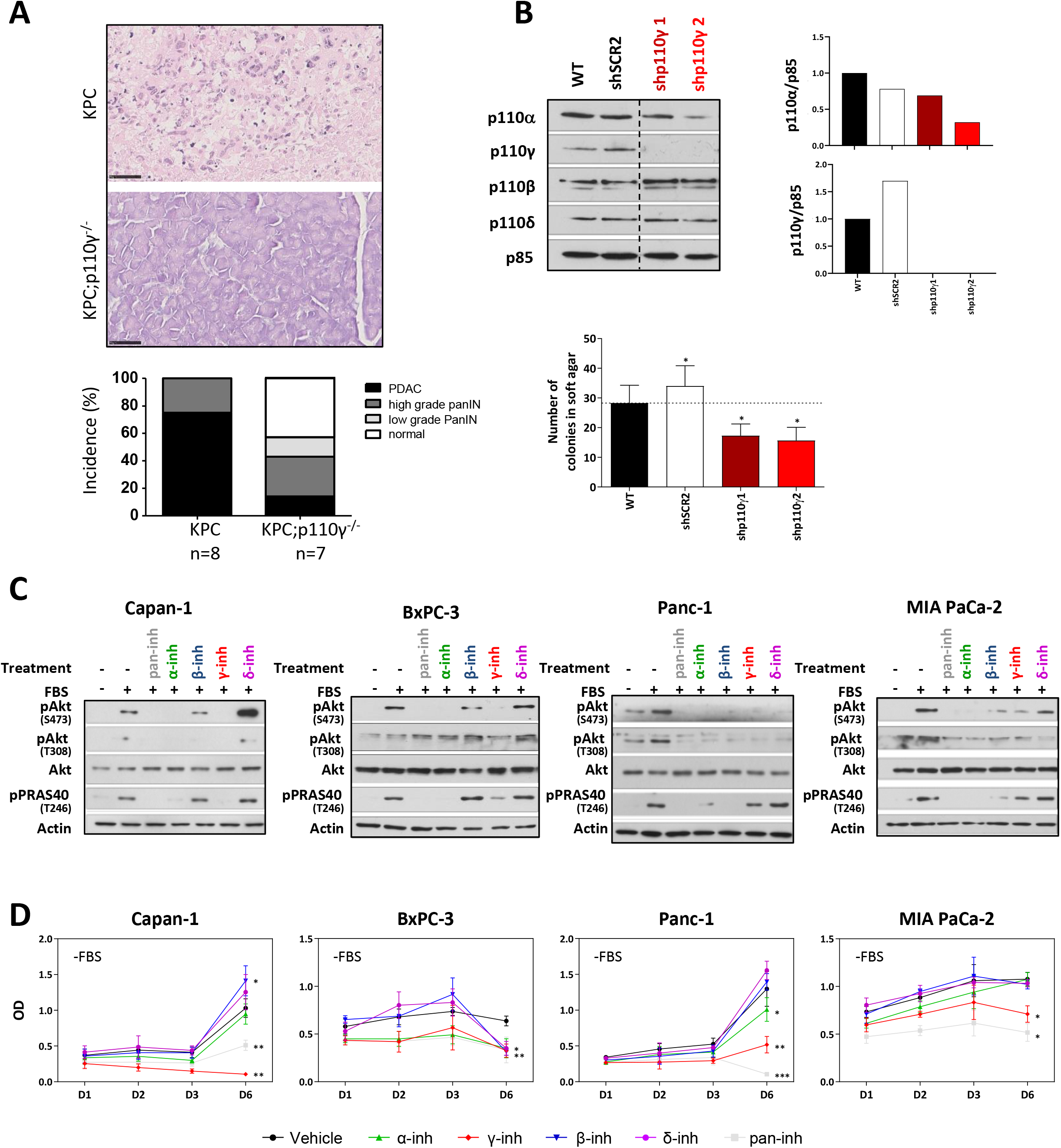
PI3Kγ-selective inhibitor sensitivity on the survival of pancreatic cancer cells. (A) Incidence of cancer in KPC (n=8 mice) versus KPC;p110γ−/− mice (n=7 mice). Representative pictures are shown. Scale=50μm (B) Validation of efficient down-regulation protein expression of p110γ targeting by genetic inhibition (shRNA) of Capan-1 cells. Number of colonies in soft agar assay of established cell lines (n=4 independent biological replicates; mean of 4 replicates ± SEM is shown). (C,D) Human pancreatic cancer cells activation of PI3K and survival were evaluated by western blot(10min) and MTT after 24h, 48h, 72h, day 6 with or without PI3K inhibitors or vehicle without 2% of FBS (n≥3 independent biological replicates; OD mean ± SEM is shown).

We tested isoform-selective inhibitors on a panel of Human pancreatic cancer cells lines. γ-inh significantly decreased pAkt at 10min in the four cell lines (**Figure 4C**, **Supplementary Figure 4A**). These experiments aimed to validate the pathway analysis on inhibitor-selective phosphoproteome, by comparing with experimental cellular outputs.

β-inh led to increased cell numbers in Capan-1 cells only (**Figure 4D**). BxPC-3 cells (non-mutated *KRAS* cell line) was sensitive to all tested inhibitors (**Figure 4D**). Cell numbers of all human pancreatic cancer cell lines were significantly decreased in time upon γ-inh or pan-inh as compared to vehicle, regardless p110γ mRNA level of expression, while α-inh was most efficient in BxPC-3 and Panc-1 cell lines (**Figure 4D**). γ-inh was almost as effective or more effective (Capan-1) that pan-inh. This was confirmed with BrdU incorporation assay, cell cycle analysis and cleaved caspase-3 analysis (**Supplementary Figure 4B-D**). Interestingly, only γ-inh treatment induced significant increase of DEVDase activity (**Supplementary Figure 4D**), confirming the selective enrichment of the “Apoptotic cleavage of cellular proteins” signaling pathway by γ-inh (**Figure 2C** **right**).

### Long-term inhibition of PI3Kγ allows compensation between PI3K isoforms in pancreatic cancer

We next analyzed the activation of PI3K canonical pathway at 24h. In Capan-1 cells, 24h inhibition with pan-inh or γ-inh led to a re-activation of p-Akt (**Figure 5A**). α-inh treatment did not show such upregulation of pAkt, in the four tested cell lines (**Figure 5A**). The combination of α-inh and γ-inh prevented this reactivation at 24h compared to γ-inh treatment. The re-activation of pAkt upon γ-inh or pan-inh treatment was observed in 3 and 2 out of 4 cell lines, respectively (**Figure 5A**, **Supplementary Figure 4A**).

**Figure 5:**
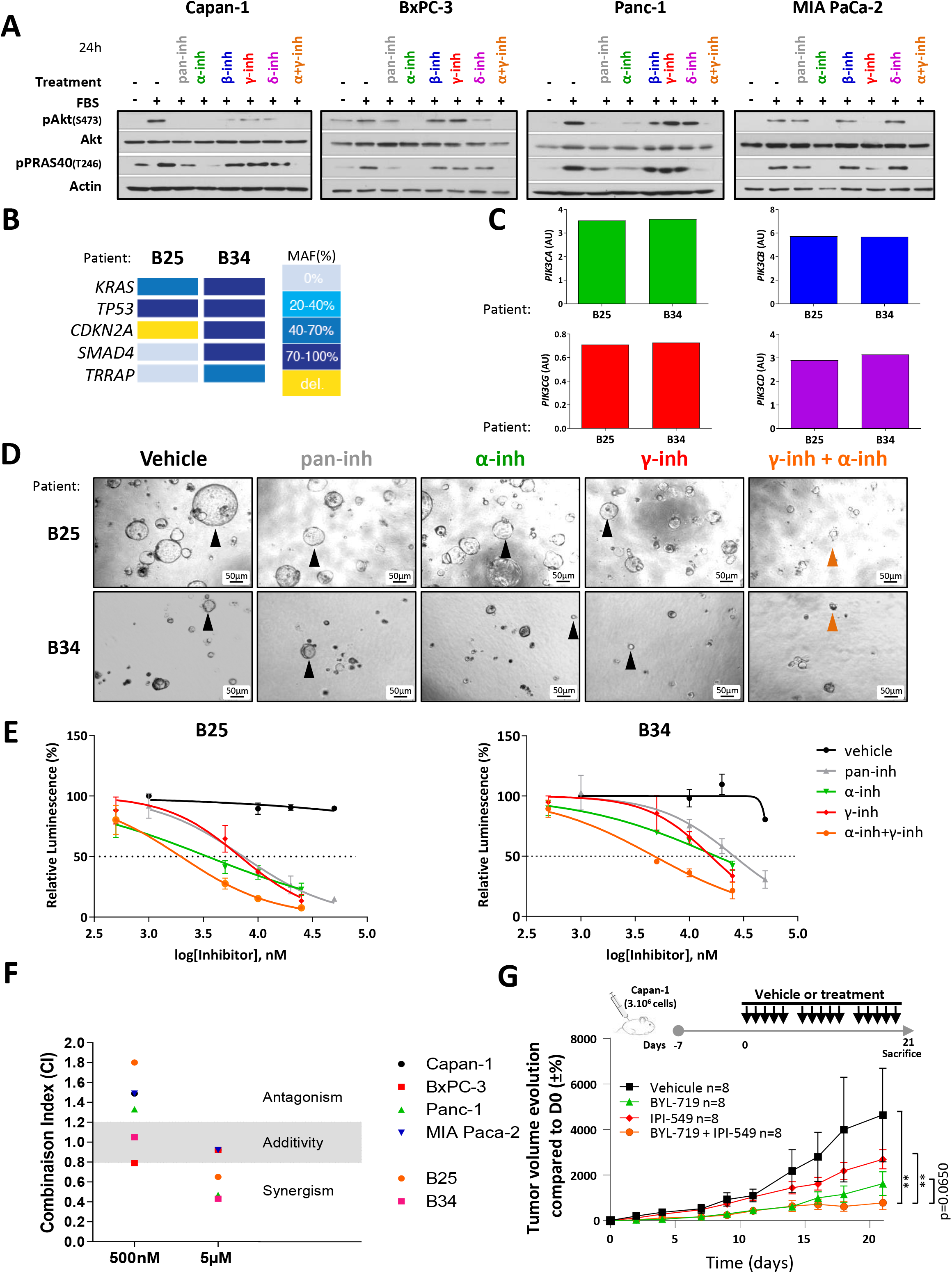
Combination of PI3Kγ and PI3Kα inhibitors lead to synergistic effect on cell survival in pancreatic ductal adenocarcinoma patients. (A) 24h of PI3K/Akt/mTOR signalling pathway activation with 2% FBS (n>3) and pan-PI3K inhibitor (n>3), PI3Kγ-inhibitor (n>3), PI3Kα-inhibitor (n>3) or dual inhibition PI3Kγ/PI3Kα (n=2) by western blot using indicated antibodies in Capan-1, BxPC-3, Panc-1 and MIA PaCa-2 cells. (B) Genetic landscape of PDAC organoids (n=2 patients). MAF=Minor Allele Frequency. (C) Level of expression of PI3K-related genes (Illumina) (n=2 patients). (D) Sensitivity to PI3K inhibition, as indicated, on morphological and number of two human pancreatic cancer primary cultures cultured in organoid condition, named B25 and B34, after 5 days of treatment. Representative pictures are shown. Scale bars=50μm (n=3 independent biological replicates). Black triangle= normal organoids; orange triangle=severely altered organoids. (E) Dose-response of two human pancreatic primary organoids viability were evaluated by CellGlo Titer assay after 5 days in complete medium with or without PI3K inhibitors, as indicated (n=3 independent biological replicates; mean ± SEM is shown). (F) Evaluation of synergistic action on survival of 7 human pancreatic cancer cells cultured in 2D (Capan-1, BxPC-3, Panc-1 and MIA PaCa-2) and 3D (B25 and B34) was performed determined using CompuSyn software based on the quantitative analysis of dose–effect relationships on multiple drugs. Combinational index (CI) values were calculated to evaluate synergy. CI < 1 indicates synergistic effects, 0.8 ≥ CI ≤ 1.2 indicates an additive effect, and CI > 1 represents an antagonistic effect (n≥3 independent biological replicates; mean ± SEM was used). (G) Xenografted 7-day Capan-1 tumors were treated with vehicle, BYL-719 (25 mg/kg), IPI-549 (7.5 mg/kg) or a combination of both treatments. n=8 mice per group, experiment performed in two separate lots. Tumor volume evolution measured with caliper is represented and corresponds to the percentage evolution of tumor growth from the first day of treatment. Mean ± SEM is shown. **, p<0.01.

Interestingly, if we were able to identify selectively changed phosphopeptides in all conditions tested, pan-inh or γ-inh conditions also led to the identification of higher numbers of selective phosphoproteins (**Figure 2B**), that are known to be inter-regulated (**Supplementary Figure 3B, 3C, 3F**), and leading similar pathway enrichments (**Figure 2C**). This could also correspond to the induction of negative feedback loops. Indeed, Kinase Enrichment Analysis (KEA) showed that pan-inh and γ-inh treatment induced the phosphorylation of peptides that corresponds to RPS6KB2 (S6K2, mTORC1 downstream effector) kinase motif (p<0.05) and to MAPK9 with lower statistical power (**Supplementary Figure 5**). These bioinformatic data are line with the observed re-induction of pAkt upon pan-inh or γ-inh treatments (**Figure 5A**).

Analysis of phosphopeptides regulated selectively by PI3Kα inhibitor at 24h in **Figure 2C** and in **Supplementary Figure 3D** showed that PI3Kα regulated additional functions and cellular components compared to panPI3K/mTOR or PI3Kγ-selective phosphopeptides. We thus tested the combination of α-inh and γ-inh on cell survival and showed that this combination led to a higher efficiency than each inhibitor alone or than PI3K/mTOR inhibitor in three out of four cell lines (**Supplementary Figure 6**). The only cell line (MIA Paca-2) where the combination was not more effective on cell survival was the one where we did not detect a feedback loop on pAkt upon PI3K inhibition. Combination of γ-inh with other isoform PI3Kβ or PI3Kδ inhibitors appeared to be less synergistic (**Supplementary Figure 6**). This corroborates the phosphoproteomic analysis that identifies feedback mechanisms on S6K upon PI3Kγ inhibition.

We next asked the question if the phosphopeptides selectively regulated at long-term by either PI3Kα or PI3Kγ inhibitors correlate with the effect of each PI3K inhibitor. The changes in ratios of pGIGYF2 (T382) correlated with PI3K action on pancreatic cell survival/proliferation (**Supplementary Figure 7A**, WO2019101871-A1). Interestingly, changes of phosphorylation levels of known targets of PI3K as assessed by WB (**Supplementary Figure 4A**) did not correlate with the cellular action in the four pancreatic cancer cell lines (**Supplementary Figure 7B**).

### The combination of PI3Kα and PI3Kγ selective inhibitors is synergistic in PDAC patients

Finally, we aimed to test this synergy in a clinically relevant setting, in two pancreatic cancer patient-derived organoids (PDOs) (B25 and B34) that displayed representative genetic alterations and p110s expression (**Figure 5B, 5C**). Here, we observed that the equimolar combination of PI3Kα-inh and PI3Kγ-inh led to an increased sensitivity to PI3K inhibition as assessed with the metabolism measurements of organoids (**Figures 5D, 5E**). In cell lines and PDOs tested, combined treatment had additive (2/6) or synergistic effect (4/6 including the two PDOs) at high concentration (5μM) (**Figure 5F**, **Supplementary Figure 8A**). At lower concentration (500nM), combination was antagonistic for the cell lines except for the non-mutant KRAS cell line BxPC-3 and the PDO B34.

For the *in vivo* proof-of-concept, we used alone or in combination two isoform-selective inhibitors that are approved for cancer treatment (*8*) tested in PDAC (*36*) or in phase II of clinical trial (https://clinicaltrials.gov/ct2/show/NCT03961698), respectively the α-selective inhibitor BYL-719/Alpelisib (25 mg/kg) and the γ-selective inhibitor IPI-549 (7.5 mg/kg). We confirmed with these inhibitors (**Supplementary Figure 8B, 8C**) the results obtained with A66 and AS-252424 in Capan-1 as shown in **Figure 4 C, 4D** and **Figure 5A**. The combination of therapeutic agents that inhibits potently PI3Kα and PI3Kγ was well tolerated and statistically more efficient that the treatment of each inhibitor alone (**Figure 5G**, **Supplementary Figure 8D**).

Overall, we observed a good correlation between the global phosphorylation-regulated pathway analysis and measured cellular and tumoral effects upon PI3K inhibitor-selective pressure, providing the first evidence of isoform-selective downstream pathways. These data also suggest that complete inhibition of PI3K with strong efficiency on PI3Kα and PI3Kγ is the most effective strategy for pancreatic cancer patients (WO2019073031-A1). This specific combination suppresses PI3K/Akt pathway in long-term manner, decreases efficiently cell survival and prevents tumor cell adaptation to PI3K signal blockage.

## Discussion

Whether critical signal nodes can be circumvented is a fundamental question in tumoral biology and therapy resistance. Using a phosphoproteomics screen, we demonstrate that each PI3K inhibitor with distinct isoform selectivity displays differential effects on the phosphoproteome of pancreatic cancer cells. Reciprocal class I PI3K signaling is the exclusive product of inhibition-imposed pressure. Accordingly, a selective and strong inhibition of PI3K isoforms is able to induce selective pathway rewiring which could facilitate selective resistance (**Graphical abstract**).

Preclinical data in pancreatic cancer have indicated for a long time that PI3K could be a suitable target in PDAC patients (initial study using Wortmannin in xenografts of Human pancreatic cancer cell lines (*37*)). There is now an urgent need to prevent feedback loops involved in pancreatic cancer progression upon pan-PI3K treatment. One strategy consists of identifying tumors which are more likely to respond. Indeed, it was recently shown that *p27kip1*, encoding a cyclin-dependent kinase inhibitor, facilitated NVP-BEZ235 (PI3K/mTOR inhibitor) sensitivity in a gene dose dependent fashion and knockdown of *p27kip1* decreased NVP-BEZ235 response in murine PDAC cell lines (*38*).

In pancreatic cancer cells, dual PI3K-mTOR inhibition induces rapid over-activation of MAPK pathway (*39*) whereas the treatment with PI3K and MAPK inhibitors were more efficient in preclinical models when used in combination than alone (*39*–*41*). Our kinase prediction data at 24h seem to indicate that both pan-PI3K and PI3Kγ-selective inhibition appear to activate MAPK9, possibly activating c-Jun pathway. Interestingly, these data are in line with our previous work where PI3K-driven NF-κB activation negatively controls JNK activation (*42*). In a future work, those combinative strategies could be tested with PI3K isoform inhibitors.

We propose that, further to MAPK signaling (*39*–*41*), adaptive response to PI3K inhibition also unleash compensatory inter-isoform selective signaling that is dependent on the isoform-selectivity of the inhibitor; this could be circumvented by well-balanced isoforms specific PI3K inhibitors. Given our results obtained *in vitro* with the first generation of isoform-selective PI3K inhibitors, we are convinced that the clinically approved Alpelisib/BYL-719 (*8*) combined with IPI-549 (*27*) that we tested in a xenograft model should be further evaluated in more complex preclinical models (e.g. KPC mice) as well as in PDAC clinical studies.

Us and others have mostly described: either 1- immediate compensations/redundancy between isoforms that are activated through similar mechanisms (e.g. PI3Kα and PI3Kδ downstream RTK (*43*); or PI3Kγ and PI3Kβ downstream GPCR (*7*)), or 2- delayed (in 1-2 week time) compensation/redundancy between the two ubiquitously expressed PI3K, PI3Kα and PI3Kβ, via genetic induction of their selective of mode activation (e.g. overexpression of RTK, mutation of PTEN) (*11*, *12*, *44*). We identify here a possible novel mode of resistance, which is based on rapid rewiring network. Next study will need to confirm that it is isoform-selective and not inhibitor-selective as well as dissect the possible mechanisms of signal rewiring.

Given the fact that PI3Kγ and PI3Kβ isoforms are both downstream of GPCRs (*7*), we were surprised to observe that the inactivation of these two isoforms did not lead to similar phosphoproteomic profiles, nor to similar alteration of cellular functions. One explanation of the poor effect of TGX-221 on these cells despite the observed reduced p-Akt levels could be the concomitant increased number of significantly increased p-peptides.

Of note, based on our results, isoform cross-compensation (in a context of non-mutated PI3K or of no loss of PTEN and of KRAS-mutant or not) appears to involve PI3Kα and the lowly expressed PI3Kγ. Because low level of expression of p110s could be difficult to detect, our data show the importance of determining the level of activation of each class I PI3K possibly with isoform selective gene signatures representative of their activity (*45*). It is also surprising that despite PI3Kγ having low levels of expression, its selective inhibition in serum-deprived condition was effective in all tested cell lines as others reported (*26*, *46*). Importantly, the level of an identified PI3Kγ-selective phosphopeptide (phopho-GIGYG2 T382) was correlated with PI3K sensitivity. We hence propose that the level of phosphorylation of this selective target could be tested as a predictive marker of sensitivity, attesting the induction of selective compensation and efficiency of combinatory treatment with PI3Kα and PI3Kγ inhibitors.

Specific engagement of either PI3Kα or PI3Kγ downstream different types of Kras mutations was recently described (*18*, *19*). It is also interesting to note that the synergistic effect of the tested combination was found in the mutated Kras cell lines, and that the PDO with increased frequence of Kras allele mutations (B34) displayed an increased combination index. These intriguing data prompt the better delineation of PI3Kα and PI3Kγ possible cooperation in various Kras-mutant contexts.

So far, only multi-combinatorial therapies displayed positive clinical outcome for PDAC patients (*22*). Isoform-specific drugs are expected to induce fewer secondary effects (*3*) and could thus be included in these multi-drug combinatorial therapies. As we show that selective inhibitors at high doses induce selective feedback and resistance mechanism -, these can be counteracted with the increasing arsenal of PI3K isoform-selective agents that are available for clinical use, in particular with Alpelisib/BYL-719 and IPI-549 (*8*, *27*). In sum, results shown here highlight that defining pharmacological profiles that are well-balanced towards each class I PI3K isoforms is key to therapeutic success.

## Material and Methods

### Reagents

Reagents were purchased as follows: for *in vitro* assays, pan-PI3K and isoform-selective PI3K inhibitors from Axon Medchem; Gemcitabine was a kind gift of hospital (IUCT-O, France); MTT was from Euromedex (4022). All PI3K inhibitors were resuspended in DMSO, corresponding at the vehicle condition. All products were resuspended according to the supplier's instructions.

### Cell lines and tissue samples

Human pancreatic cell lines (Capan-1, BxPC-3, PANC-1, MIA PaCa-2) came from American Type Culture Collection (ATCC), human acute myeloid leukemia cell line (MOLM4) and murine pancreatic cancer cell lines (DT4994, DT6585, DT6606, DT8442, DT8661, R221, R259) were made in house or a kind gift from Dieter Saur (Klinikum rechts der Isar der TU München, Germany); for validation of genotype (*16*). Absence of mycoplasma contamination was verified periodically by PCR and maintained in culture for a maximum of 15 passages after thawing. Capan-1, PANC-1 and MIA PaCa-2 cells were authenticated (STR method, Eurofins). Pancreatic cancer patient-derived organoids were either derived from surgical resection (B25) or endoscopic fine needle aspiration (B34) at Klinikum rechts der Isar der TU Mu◻nchen, Germany. Patients were enrolled and consented in writing according to the institutional review board (IRB) approval project-number 207/15 of the Technical University Munich. The studies were conducted in accordance with the Declaration of Helsinki. PDOs were characterized using whole exome sequencing and RNAseq as described previously (*47*). Human normal and adenocarcinoma pancreatic samples (>30% tumoral cells) were selected at IUCT-O clinic, and collected according French and European legislation (CRB Biobank, France with following ethical authorization numbers BB-0033-00014, DC-2008-463, AC-2013-1955). KPC and KPC; p110γ−/− mice were obtained and their genotype verified as described in (*27*).

### *In vitro* culture of pancreatic cell lines and cell assays

Human pancreatic cancer cell lines Capan-1 and BxPC-3 were cultured in RMPI 1640 medium. PANC-1, MIA PaCa-2 and all murine pancreatic cancer cells were cultured in Dulbecco’s Modified Eagle’s Medium with 4.5g of glucose (D6429, Sigma). All media were supplemented with 10% fetal bovine serum (Eurobio), 1% glutamine (G7513, Sigma) and 1% antibiotics (penicillin/streptomycin, P0781, Sigma). Patient’s organoids cell lines were cultured in special medium (Reichert composition,(*47*)). Specific methods to analyze cell proliferation/survival, mRNA and protein expression levels are described in **Supplementary Material &Methods**.

### SILAC phosphoproteome and Bioinformatics analysis

Light amino acid-labelled and heavy amino acid-labelled Capan-1 cells (respectively called thereafter “light” and “heavy” cells) were cultured as described in **Supplementary Material &Methods.** For each biological replicate, light lysates were mixed with the same heavy lysates at a 1:1 ratio for a total amount of 6 mg (**Figure 1C**). Protein samples were reduced with 100 mM DTT (Sigma, D9163) for 35 min at 57°C and then handled according to the FASP (Filter Aided Sample Preparation) digestion protocol (*48*) using Amicon Ultra-15 Centrifugal Filter device (10kDa cut-off, MILLIPORE, UFC901096). Protocol for the enrichment with TiO_2_ beads is based on Larsen et al., (*49*) and Jensen et al. (*49*). SILAC samples (TiO_2_ enriched peptides) were resuspended with 2% acetonitrile, 0.05% TFA and analyzed by nano-LC-MS/MS using an UltiMate 3000 system (Dionex) coupled to LTQ-Orbitrap Velos mass spectrometers (Thermo Fisher Scientific, Bremen, Germany). All 40 mass spectrometry proteomic files have been deposited to ProteomeXchange Consortium with the dataset identifier as listed in **Supplementary Table 1**. For peptide identification, raw data files were processed in Proteome Discover 1.4.1.14 (Thermo Scientific) and searched against SwissProt human fasta database of Mascot (2014-06, sprot_20140428.fasta, 542782 sequences, high and medium confidence, Q-value = 0.5-0.1). Peptides were further filtered using Mascot significance threshold S/N = 1.5 and a FDR <0.01 based on q-Value (Percolator). Phospho-site localization probabilities were calculated with *phospho*RS 3.1 (maximum PTMs per peptide 10, maximum position isoforms 200). Phosphopeptides filtered with Proteome Discoverer 1.4.1.14 (see criteria in **Supplementary Material &Methods**) were isolated from peptides. Only the ratios which were changed above and below the thresholds were processed for further analysis as described in **Supplementary Material &Methods**.

### *In vivo* experiment

All animal procedures were conducted in compliance with the Ethics Committee pursuant to European legislation translated into French Law as Décret 2013-118 dated 1st of February 2013 (APAFIS 3601-2015121622062840).

Capan-1 cells were tested for their absence of mycoplasma infection prior amplification and injection. 3×10^6^ exponentially growing Capan-1 cells were subcutaneously in Nude/balbc Mice (Charles River, 9 weeks old, females). After 1 week implantation, we gavaged the mice 5 day a week with vehicle (0.5% methyl cellulose with 0.2% Tween-80) or with BYL-719/Alpelisib (25 mg/kg) (MedChemExpress) (*50*), IPI-549 (7.5 mg/kg) (Biorbyt) (*51*) alone or in combination. Toxicity parameters were assessed longitudinally with follow-up of mice weight, glycemia, blood cell counts (using Yumizen H500 hematology analyzer (HORIBA)). The drug dosage with treatment 5 day a week lasted for 3 weeks, time at which the vehicle group had to be euthanized for ethical reasons. Tumor volume was measured with a caliper and calculated using the formula V= (4/3) x π x (Length/2)2 x (width/2).

### Statistics

Statistically significant differences were performed with GraphPad Prism using the T-tests (paired test): * P < 0.05, ** P < 0.01, *** P < 0.001. Non-significant (ns) if P > 0.05. For *in vivo* experiments, Mann-Whitney test was used: ** p < 0.01, p value is indicated for nearly significant values.

## Acknowledgments

We thank members of SigDYN group for constructive discussions, as well as members of CRCT core platforms, vectorology, cytometry and imaging, B Couderc for p110s-shRNA plasmids.

## Author contributions

CC, DT, DZ, TN, CC, RDF, TB, CP: experiments; CC, DT, MBE, BMP, JGG: formal analysis of data; CC, DT, MBE, BMP, PF, GSB, BSO, MR, JGG: methodology; CC, JGG: visualization of data; CC, TD, BMP: material and methods writing; CC, JGG: review of writing; all authors: editing MS; PC, HE, BGA, MR: provided samples; BSO, MR, JGG: supervision; JGG: conceptualization, funding acquisition, project administration, project supervision, validation, writing – original draft.

## Data and materials availability

Phosphoproteomics data has been deposited in Pride / ProteomeXchange with the dataset identifier PDX008410 the 11/12/2017. PDO, plasmids for sh expression used in the manuscript require an MTA.

---

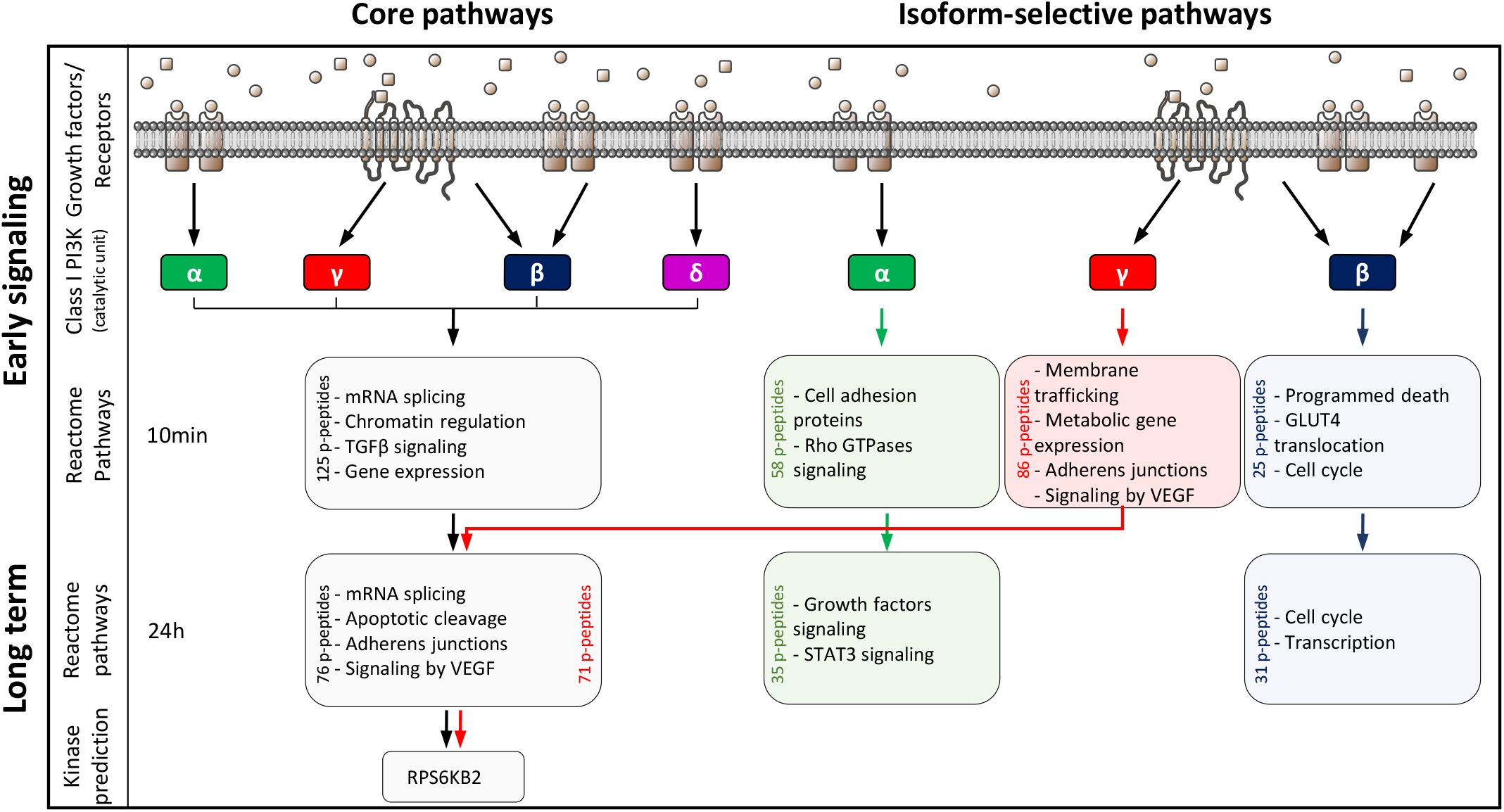

