## Supplementary material for "Phosphoproteomics identifies PI3K inhibitor-selective adaptive responses in pancreatic cancer cell therapy and resistance": Figure S1 to S8

### Supplementary Figure 1

**A**

| PDAC Patient | Gender | Diagnosis age | Survival (months) | Treatment |
|---|---|---|---|---|
| **1** | F | 60 | 32 | Gemzar (6 months) + Folfirinox (12 cycles) |
| **2** | M | 61 | 43 | Gemzar (6 months) + Xeloda + Folfirinox and Folfox (12 cycles) + Folfirinox (6 cycles) |
| **3** | M | 56 | MD | Gemzar (6 months) |
| **4** | M | 76 | MD | Gemzar (6 months) + Radiotherapy |
| **5** | F | 57 | 24 | Gemzar (6 months) + Folfirinox (2 cycles) + Xeloda + Radiotherapy |
| **6** | M | 50 | MD | Gemzar (6 months + Folfirinox (18 cures) + Radiotherapy |
| **7** | F | 69 | MD | Gemzar (6 months + Folfirinox (6 cures) |
| **8** | F | 67 | 59 | Gemzar (6 months + Folfirinox (6 cures) |
| **9** | F | 85 | MD | MD |
| **10** | F | 68 | 17 | Gemzar (6 months) + Folfox (4 cures) + Folfirinox (10 cures) |
| **11** | M | 81 | MD | Gemzar (6 months) |
| **12** | M | 54 | 0 | MD |
| **13** | F | 45 | MD | Gemzar (6 months) |
| **Median** | | **60** | **28** | |
| **Mean** | | **62.7** | **29.17** | |

Missing data = MD

**B**

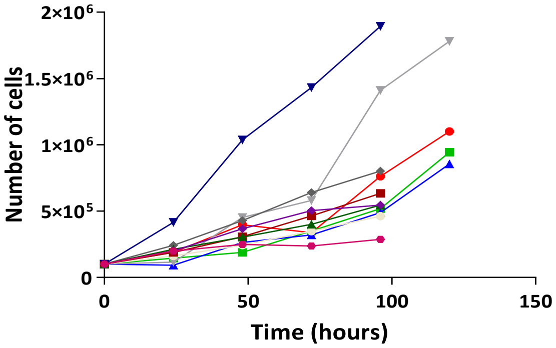


| Cell Lines | Doubling Time (hours) |
|---|---|
| Capan-1 | 34.8 |
| BxPC-3 | 37.1 |
| Panc-1 | 38.8 |
| Mia PACA-2 | 28.9 |
| DT4994 | 31.9 |
| DT6585 | 43.4 |
| DT6606 | 36.0 |
| DT8442 | 48.0 |
| DT8661 | 25.0 |
| R221 | 41.2 |
| R259 | 77.1 |

**C**

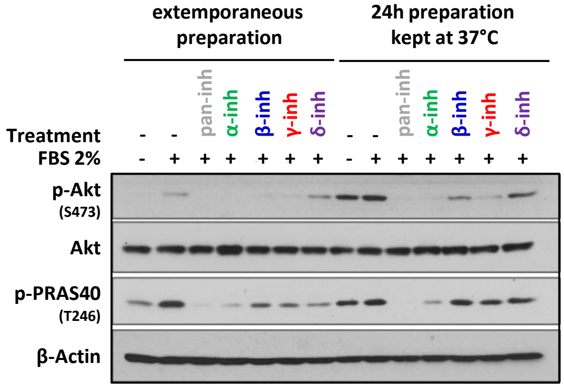

#### Supplementary Figure 2

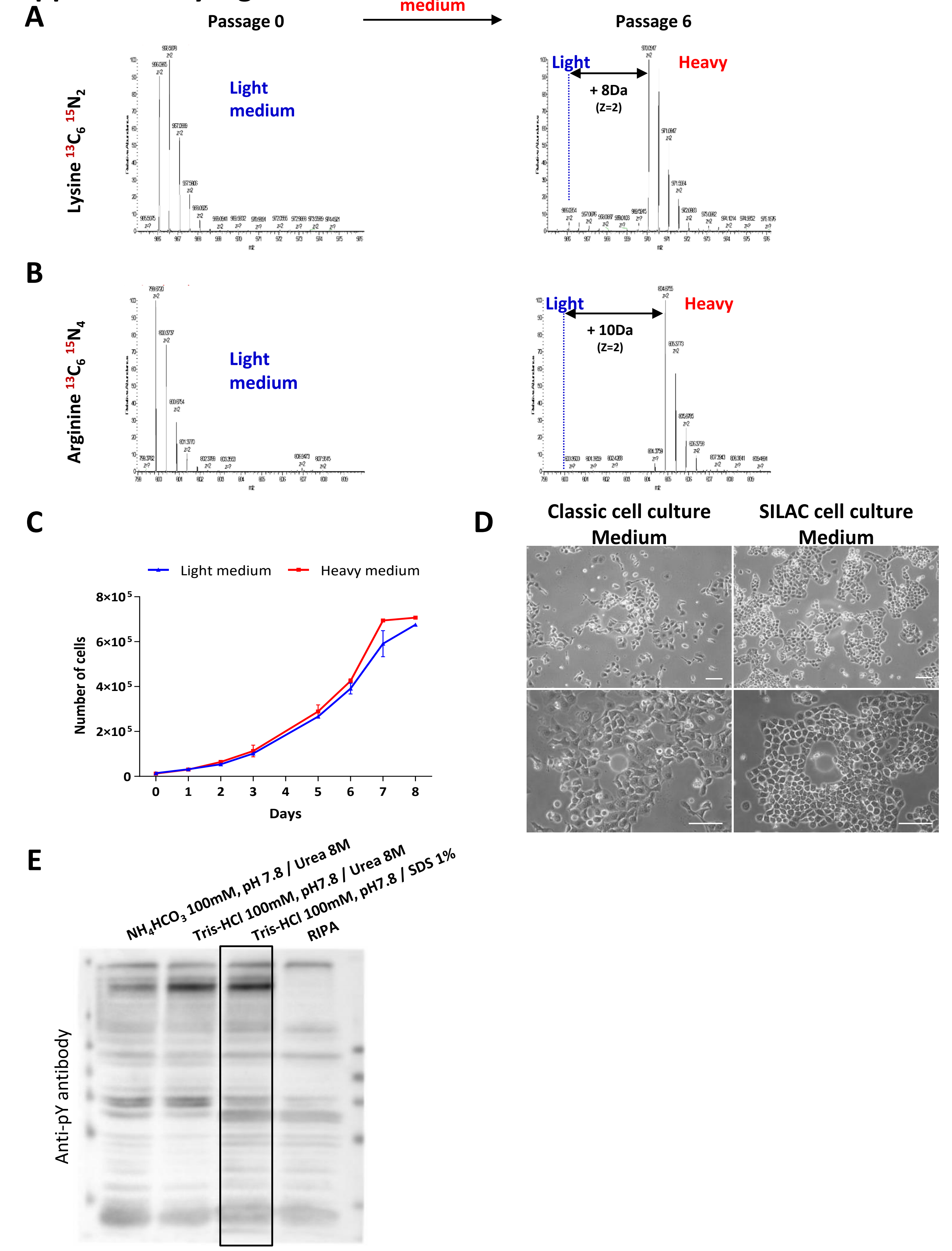

### Supplementary Figure 2

**F**

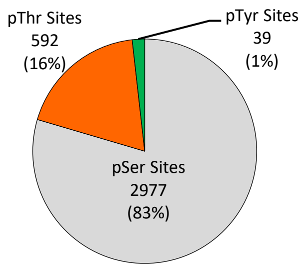


**G**

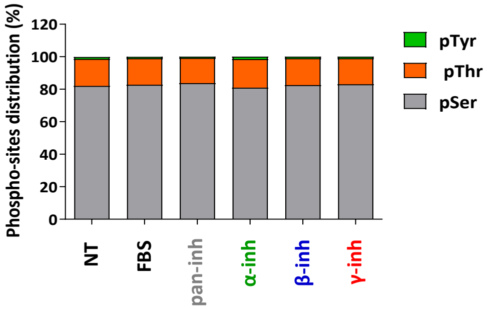


**H**

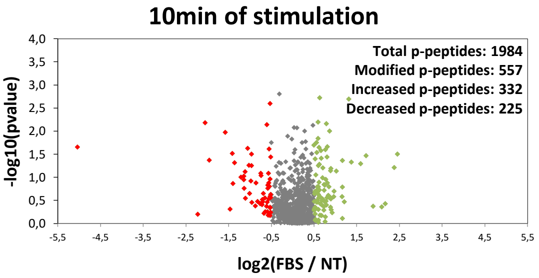


**I**

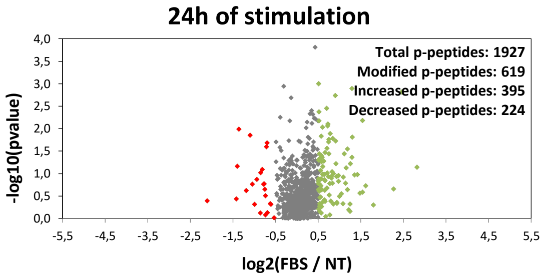


**J** **Phospho-peptide of pPRAS40 (T246): LNtSDFQk**

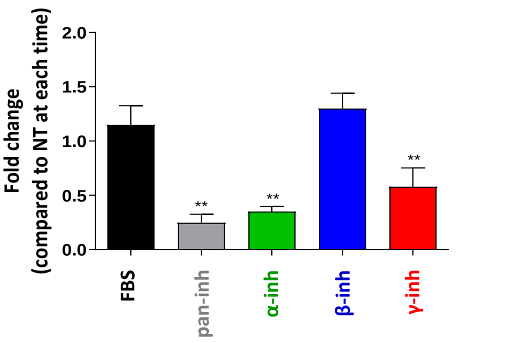

### Supplementary Figure 3

nc = no connectivity to PI3Ks

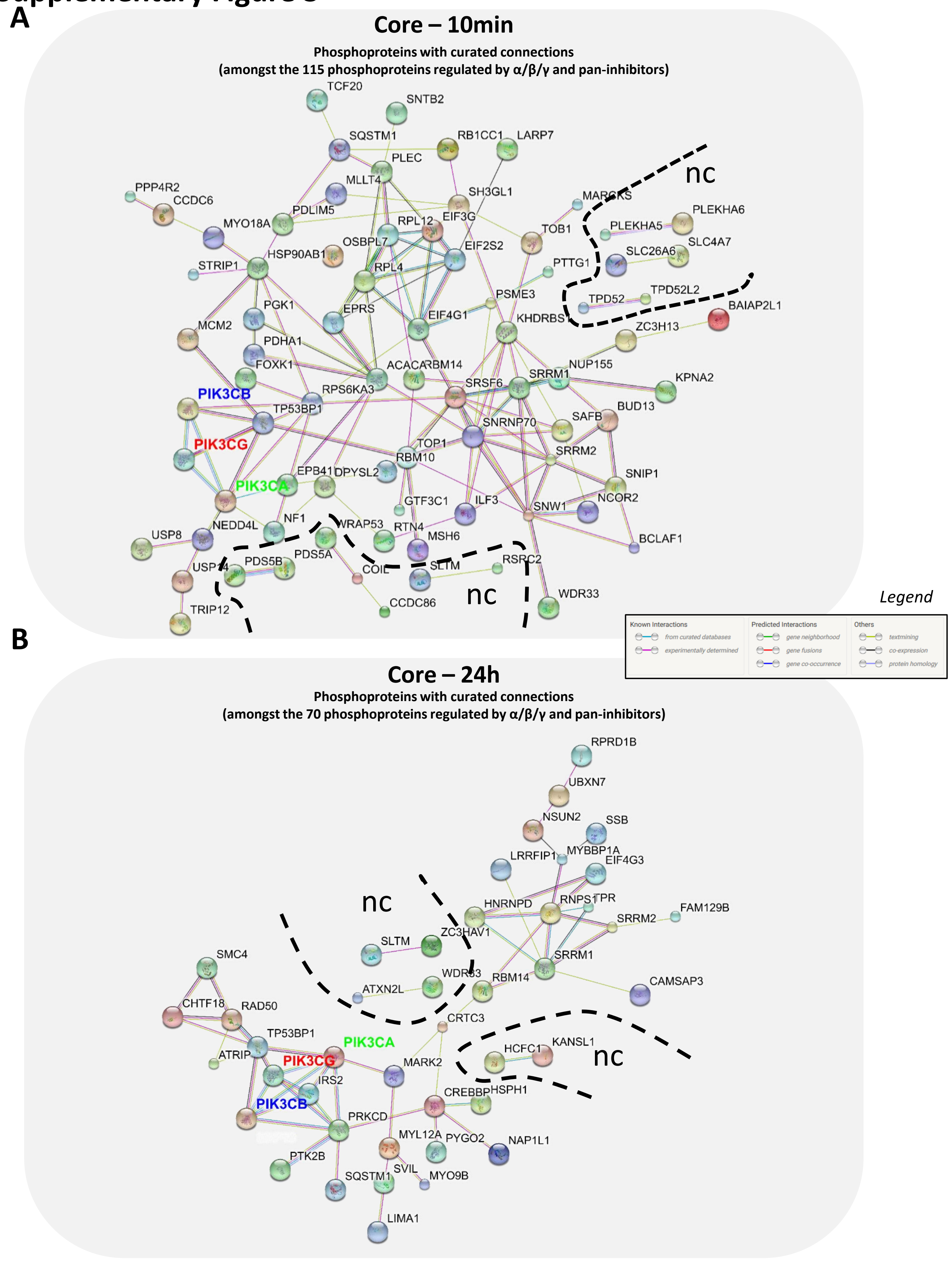

### Supplementary Figure 3

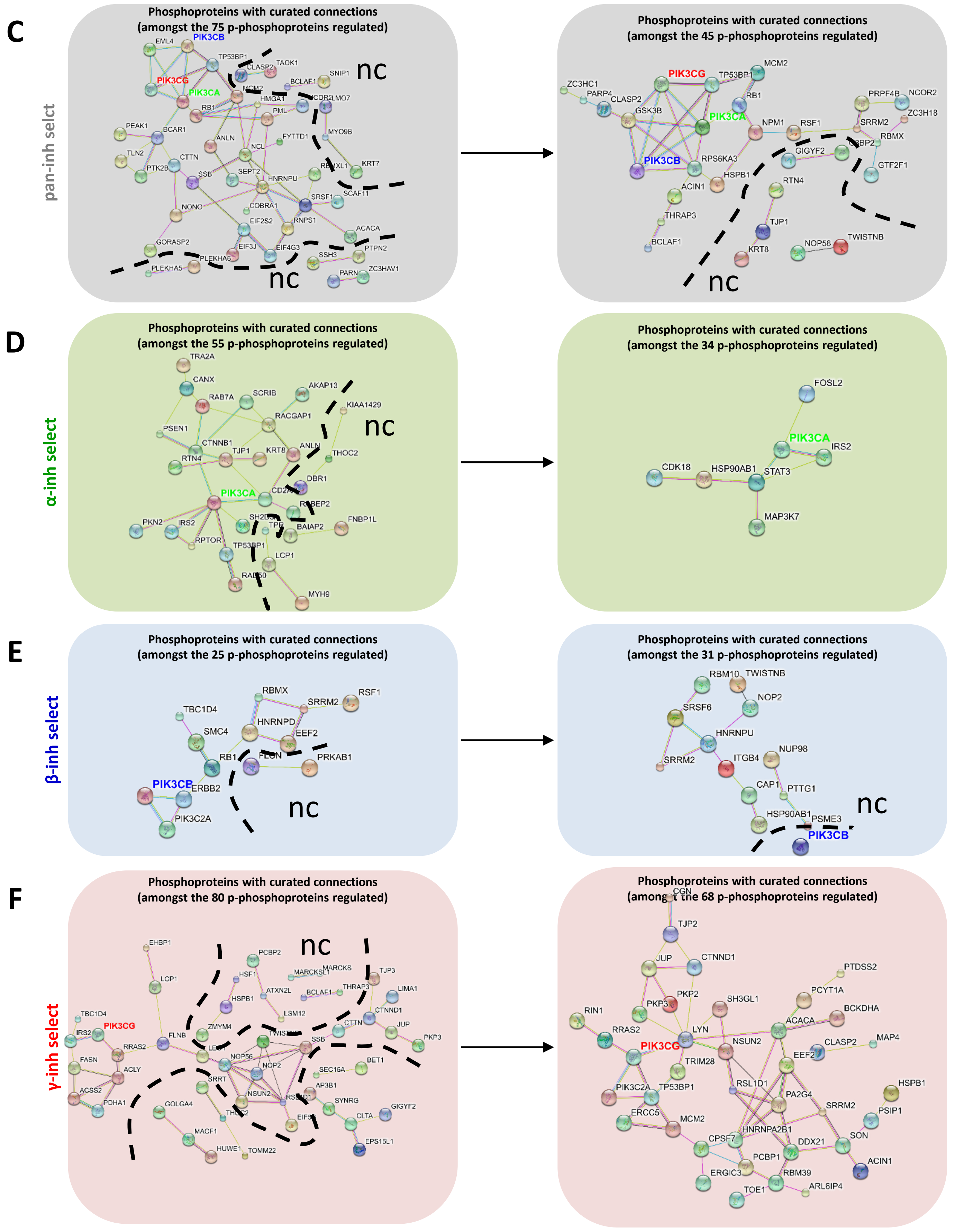

### Supplementary Figure 3

**G** Cellular Component (GO)

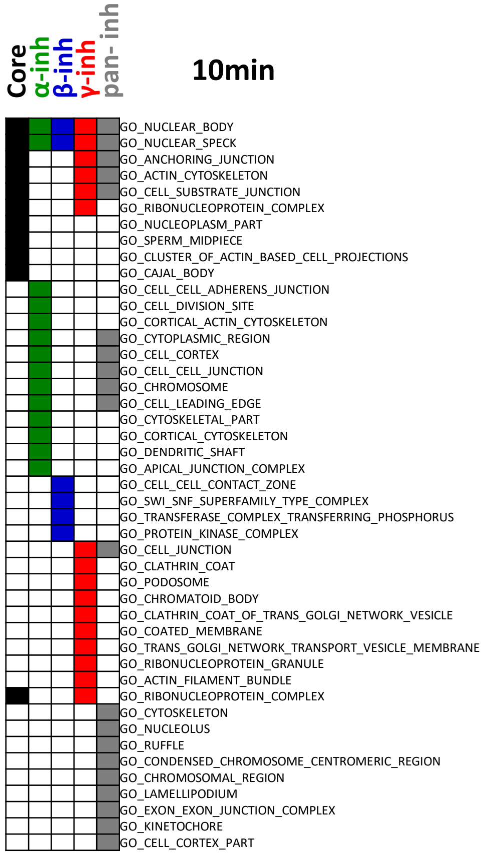


**H**

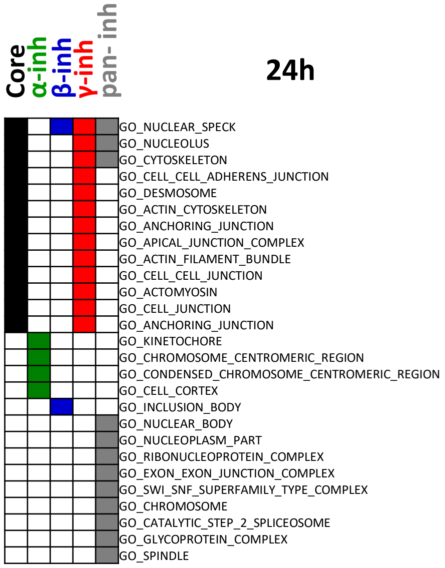


**I** Molecular Function (GO)

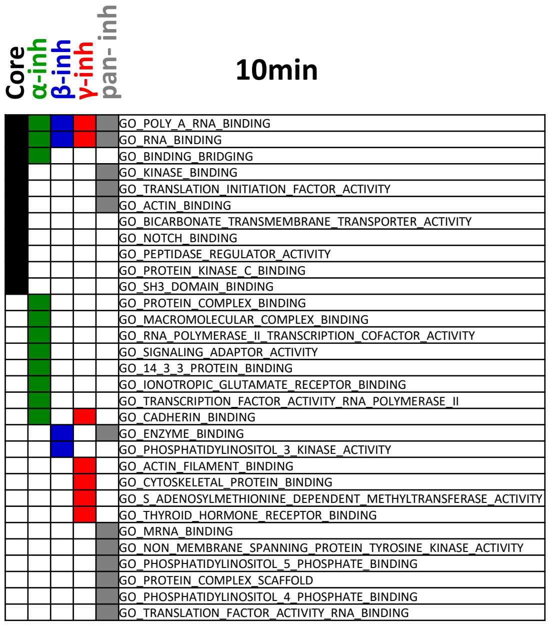


**J**

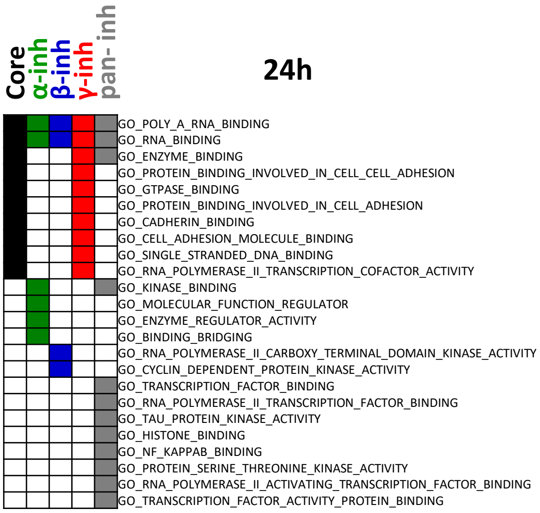

### Supplementary Figure 4

A

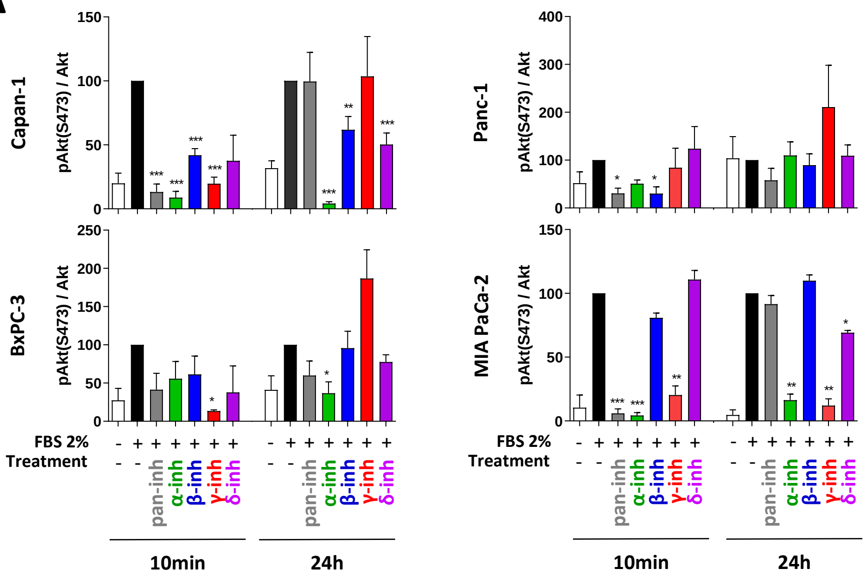


B

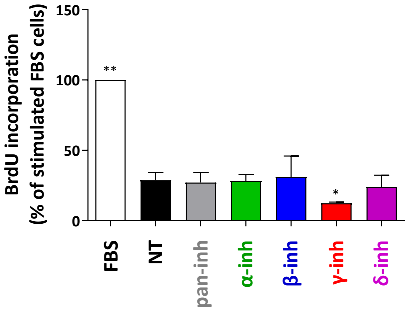


C

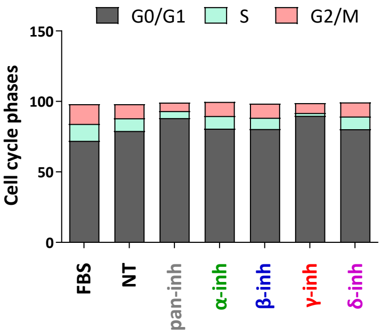


D

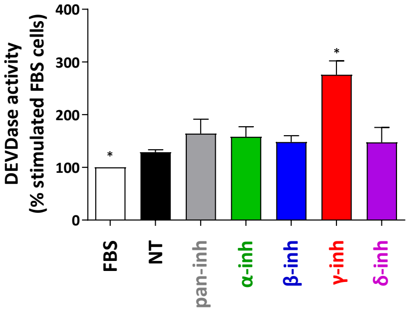

### Supplementary Figure 5

**A** 10min **B** 24h

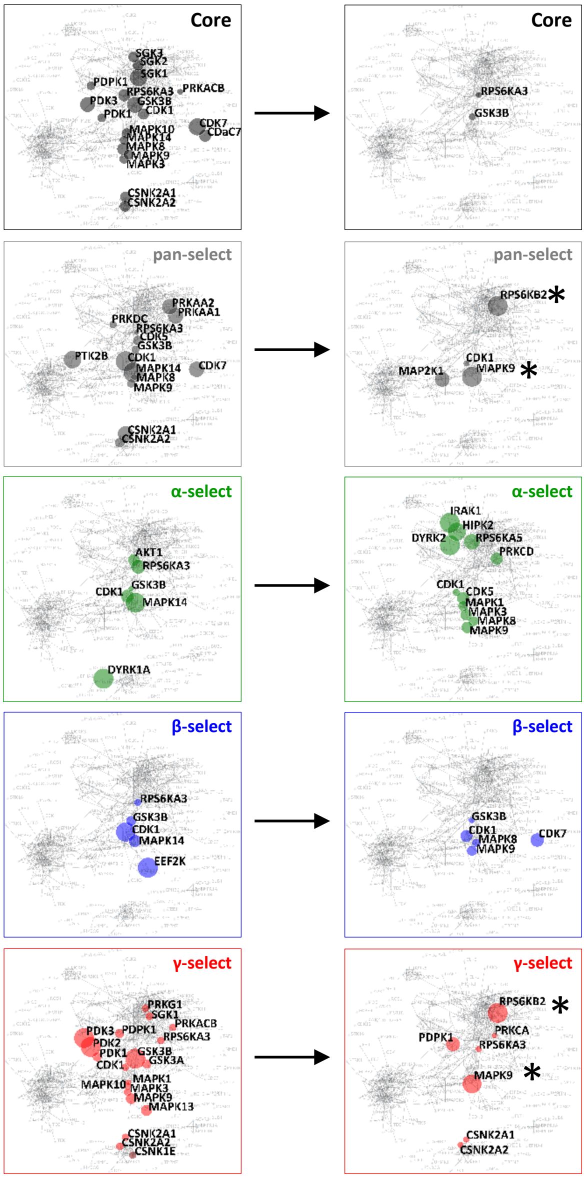


**C**

| 10min | Kinases enriched | p-value |
|---|---|---|
| Core | SGK1 | 0.0413 |
| | CDK7 | 0.0436 |
| | GSK3B | 0.0564 |
| | PDK3 | 0.0696 |
| | RPS6KA3 | 0.124 |
| | CDC7 | 0.124 |
| | MAPK8 | 0.124 |
| | CDK1 | 0.147 |
| | CSNK2A2 | 0.163 |
| | CSNK2A1 | 0.167 |
| | SGK2 | 0.167 |
| | SGK3 | 0.167 |
| | MAPK10 | 0.192 |
| | MAPK9 | 0.204 |
| | MAPK3 | 0.233 |
| | PDPK1 | 0.258 |
| | PDK1 | 0.258 |
| | MAPK14 | 0.264 |
| | PRKACB | 0.468 |
| pan- | MAPK14 | 0.0663 |
| | CDK1 | 0.0663 |
| | PTK2B | 0.0979 |
| | CSNK2A1 | 0.133 |
| | CDK7 | 0.133 |
| | PRKAA1 | 0.133 |
| | PRKAA2 | 0.133 |
| | CSNK2A2 | 0.325 |
| | CDK5 | 0.325 |
| | MAPK9 | 0.396 |
| | GSK3B | 0.411 |
| | PRKDC | 0.455 |
| | MAPK8 | 0.554 |
| | RPS6KA3 | 0.646 |
| α-select | MAPK14 | 0.0399 |
| | DYRK1A | 0.0399 |
| | RPS6KA3 | 0.117 |
| | CDK1 | 0.144 |
| | AKT1 | 0.185 |
| | GSK3B | 0.555 |
| β-select | EEF2K | 0.0191 |
| | CDK1 | 0.0215 |
| | MAPK14 | 0.110 |
| | GSK3B | 0.185 |
| | RPS6KA3 | 0.333 |
| γ-select | GSK3B | 0.100 |
| | PDK3 | 0.100 |
| | PDK2 | 0.100 |
| | MAPK9 | 0.327 |
| | MAPK13 | 0.327 |
| | GSK3A | 0.416 |
| | PDPK1 | 0.416 |
| | PDK1 | 0.416 |
| | SGK1 | 0.490 |
| | CDK1 | 0.508 |
| | CSNK2A2 | 0.508 |
| | PRKG1 | 0.508 |
| | PRKACB | 0.508 |
| | CSNK2A1 | 0.508 |
| | MAPK10 | 0.514 |
| | RPS6KA3 | 0.514 |
| | CSNK1E | 0.514 |
| | MAPK3 | 0.514 |
| | MAPK1 | 0.658 |

**D**

| 24h | Kinases enriched | p-value |
|---|---|---|
| Core | GSK3B | 0.226 |
| | RPS6KA3 | 0.322 |
| pan- | RPS6KB2* | 0.023 |
| | MAPK9 | 0.023 |
| | MAP2K1 | 0.067 |
| | CDK1 | 0.446 |
| α-select | DYRK2 | 0.009 |
| | IRAK1 | 0.009 |
| | HIPK2 | 0.016 |
| | RPS6KA5 | 0.03 |
| | PRKCD | 0.07 |
| | CDK5 | 0.07 |
| | MAPK9 | 0.087 |
| | MAPK3 | 0.09 |
| | MAPK1 | 0.138 |
| | MAPK8 | 0.138 |
| | CDK1 | 0.256 |
| β-select | CDK7 | 0.033 |
| | CDK1 | 0.05 |
| | MAPK9 | 0.094 |
| | MAPK8 | 0.153 |
| γ-select | RPS6KB2* | 0.05 |
| | MAPK9 | 0.061 |
| | PDPK1 | 0.14 |
| | CSNK2A2 | 0.53 |
| | CSNK2A1 | 0.53 |
| | RPS6KA3 | 0.53 |
| | PRKCA | 0.588 |

### Supplementary Figure 6

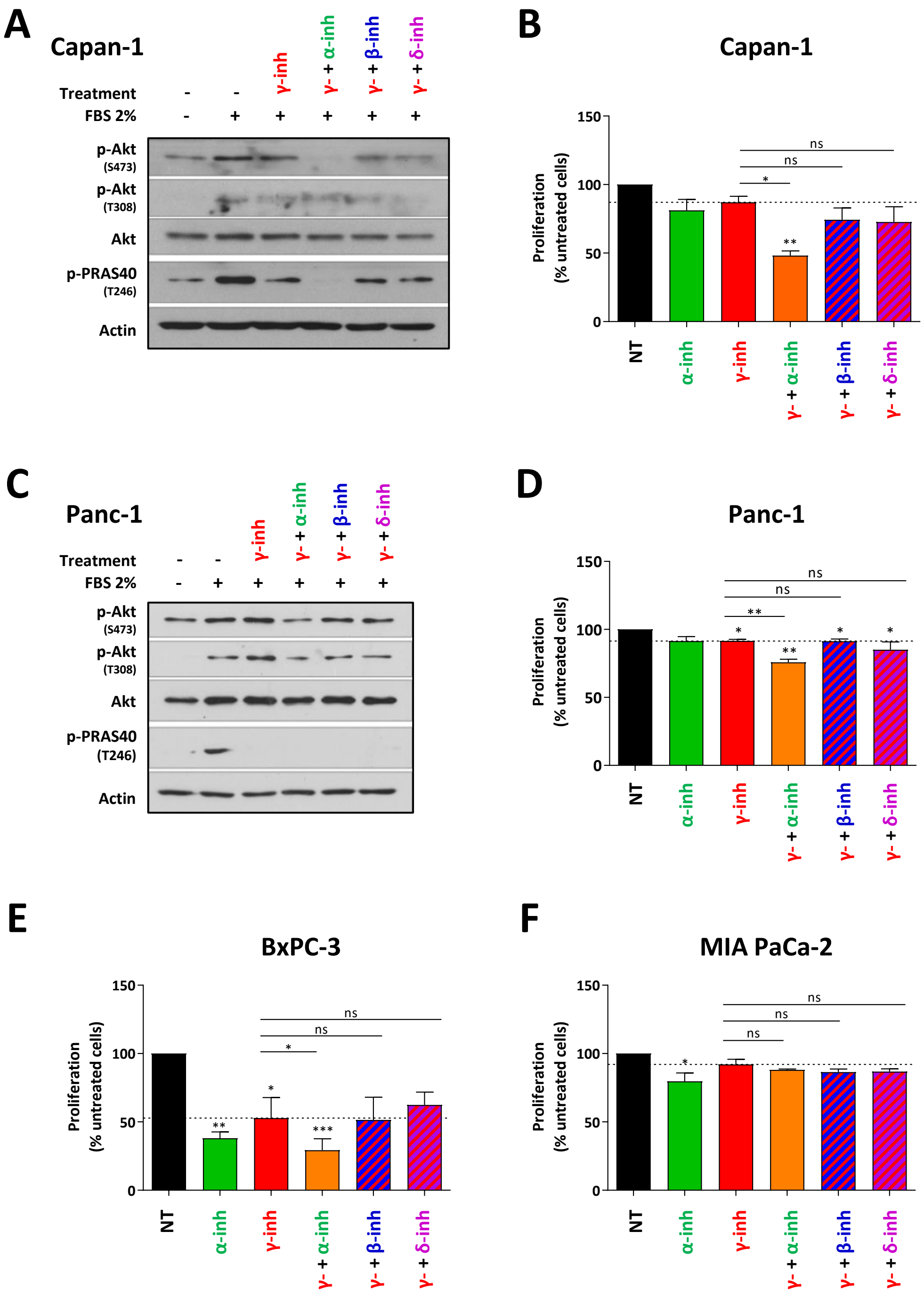

### Supplementary Figure 7

**A**

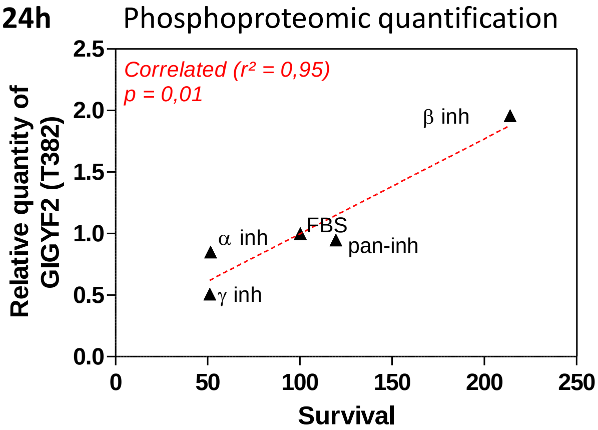


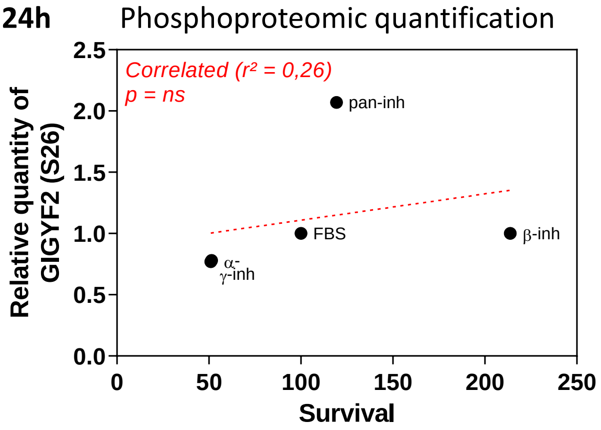


Phospho-peptide of pGIGYG2 (T382):
PGt(phospho)PSDHQSQEASQFER

Phospho-peptide of pGIGYG2 (S26):
ALSSGGSITs(phospho)PPLSPALPK

| Capan-1 | FBS | pan-inh | α-inh | β-inh | γ-inh |
|---|---|---|---|---|---|
| GIGYF2_T382 | 1 | 0.95 | 0.85 | 1.96 | 0.51 |

| Capan-1 | FBS | pan-inh | α-inh | β-inh | γ-inh |
|---|---|---|---|---|---|
| GIGYF2_S26 | 1 | 2.07 | 0.78 | 1.00 | 0.77 |

**B**

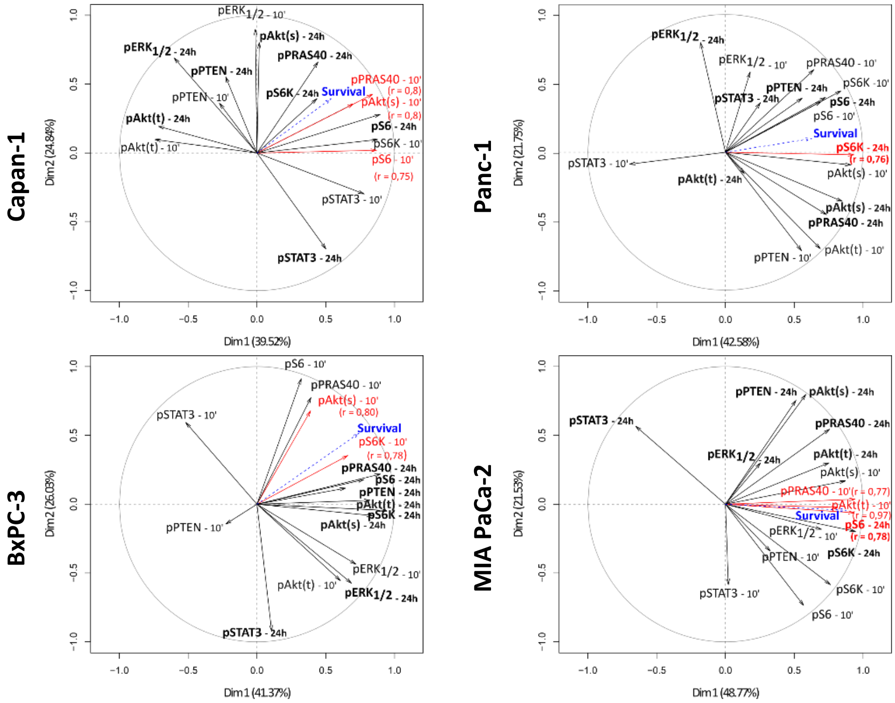

### Supplementary Figure 8

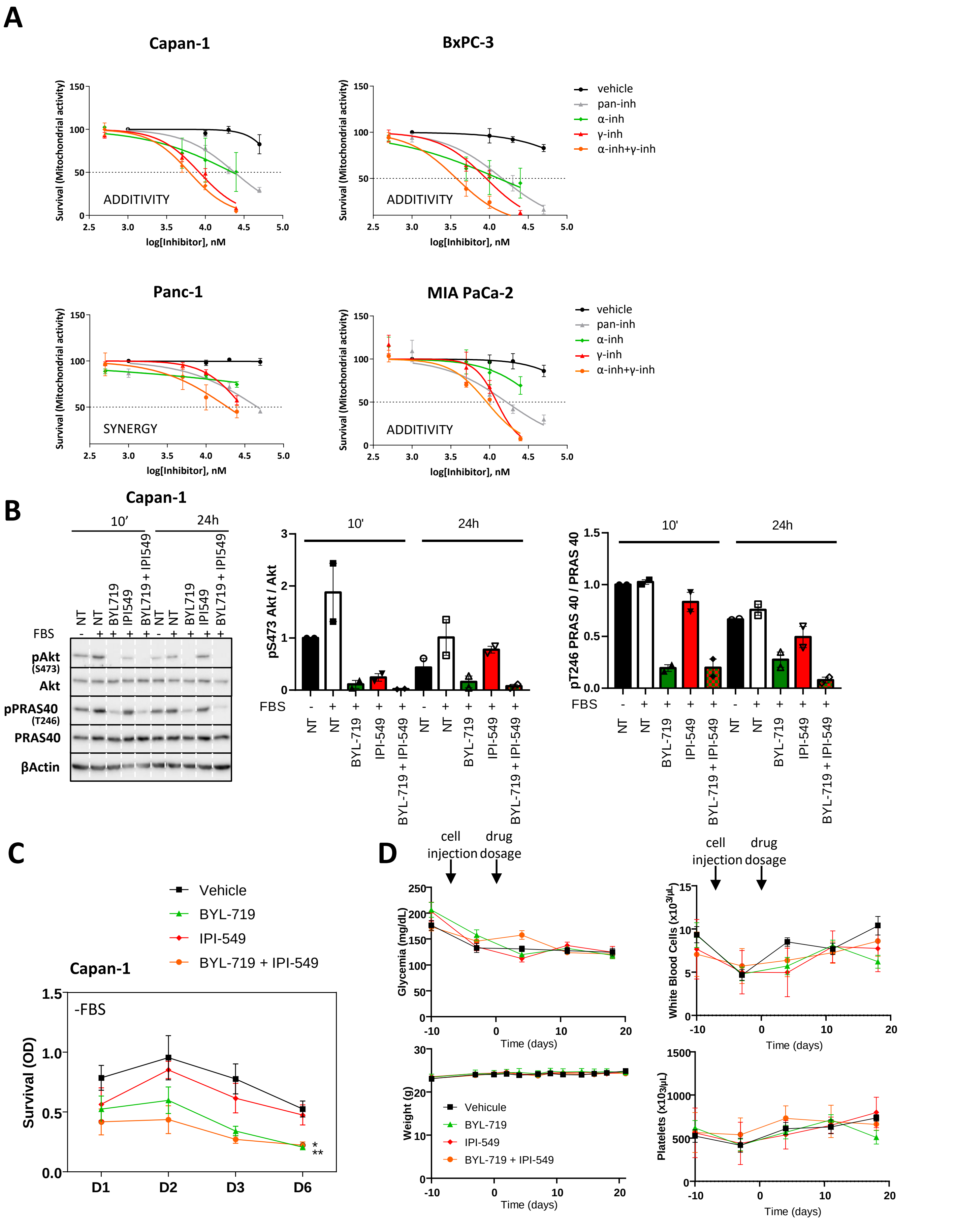
